# The *O*-Fucosyltransferase SPINDLY Attenuates Auxin-Induced Fruit Growth by Inhibiting ARF6 and ARF8 binding to Coactivator Mediator Complex in *Arabidopsis*

**DOI:** 10.1101/2024.06.26.599170

**Authors:** Yan Wang, Seamus Kelley, Rodolfo Zentella, Jianhong Hu, Hua Wei, Lei Wang, Jeffrey Shabanowitz, Donald F. Hunt, Tai-ping Sun

## Abstract

The phytohormone auxin plays a pivotal role in promoting fruit initiation and growth upon fertilization in flowering plants. Upregulation of auxin signaling by genetic mutations or exogenous auxin treatment can induce seedless fruit formation from unpollinated ovaries, termed parthenocarpy. Recent studies suggested that the class A AUXIN RESPONSE FACTOR6 (ARF6) and ARF8 in *Arabidopsis* play dual functions by first inhibiting fruit initiation when complexed with unidentified corepressor IAA protein(s) before pollination, and later promoting fruit growth after fertilization as ARF dimers. However, whether and how posttranslational modification(s) regulate ARF6- and ARF8-mediated fruit growth were unknown. In this study, we reveal that both ARF6 and ARF8 are *O*-fucosylated in their middle region (MR) by SPINDLY (SPY), a novel nucleocytoplasmic protein *O-*fucosyltransferase, which catalyzes the addition of a fucose moiety to specific Ser/Thr residues of target proteins. Epistasis, biochemical and transcriptome analyses indicated that ARF6 and ARF8 are downstream of SPY, but ARF8 plays a more predominant role in parthenocarpic fruit growth. Intriguingly, two ARF6/8-interacting proteins, the co-repressor IAA9 and MED8, a subunit of the coactivator Mediator complex, were also *O*-fucosylated by SPY. Biochemical assays demonstrated that SPY-mediated *O*-fucosylation of these proteins reduced ARF-MED8 interaction, which led to enhanced transcription repression activity of the ARF6/8-IAA9 complex but impaired transactivation activities of ARF6/8. Our study unveils the role of protein *O*-fucosylation by SPY in attenuating auxin-triggered fruit growth through modulation of activities of key transcription factors, a co-repressor and the coactivator MED complex.

## INTRODUCTION

In flowering plants, the developmental transition from ovary to fruit, termed fruit initiation or fruit set, usually occurs after fertilization. Phytohormones play key roles in controlling fruit set, growth and maturation^1–4^. Among them, auxin synthesized in developing seeds promote fruit set and subsequent growth. Treatment of the unfertilized ovary with auxin can induce parthenocarpy, i.e., seedless fruit formation without fertilization, indicating that activation of auxin signaling is essential for fruit initiation^5–7^. Parthenocarpy is often a beneficial trait in crops. Seedless fruits are preferred by the consumers, and this trait is also associated with flavor improvement and longer shelf life^8^. Moreover, it offers more consistent fruit yield in variable environmental conditions such as elevated temperatures that can severely reduce pollen viability and limit pollinator availability both of which can limit fruit production^9^. Elucidation of the regulatory mechanism of auxin-induced parthenocarpy will contribute to improving yield stability under stressful climate conditions.

The nuclear auxin signaling pathway consists of three families of major components, (1) auxin coreceptors TRANSPORT INHIBITOR 1/AUXIN SIGNALING F-BOX (TIR1/AFB), (2) transcription co-repressors Auxin/INDOLE-3-ACETIC ACID (Aux/IAA), and (3) AUXIN RESPONSE FACTOR (ARF) transcription factors^10–14^. Aux/IAA proteins (thereafter abbreviated as IAAs) function as negative regulators of auxin signaling by binding to ARFs at their target promoters. Formation of the IAA-ARF complexes leads to the recruitment of co-repressor TOPLESS (TPL), which represses transcription by preventing ARF binding to the coactivator Mediator complex^15^. TPL may also recruit the CDK8 kinase module (CKM), a repressive module of the Mediator complex, to block transcription of ARF target genes^16^. When auxin levels increase, it promotes TIR1/AFB-IAA interactions, thereby triggering degradation of IAA proteins to release ARFs to activate the downstream auxin response pathway. *AFBs*, *IAA*s and *ARF*s all belong to multi-gene families in angiosperms, whereas much reduced gene redundancy of these auxin signaling components was found in early-emerging land plants, e.g., *Physcomitrella patens* (moss) and *Marchantia polymorpha* (liverwort)^17^. IAA proteins contain three domains: Domain I includes a transcriptional repression EAR motif that recruits TPL, domain II contains an interaction motif for TIR1/AFB, and the conserved C-terminal Phox and Bem1 (PB1) domain is required for multimerization between IAAs and ARFs^10^. ARFs contain a conserved N-terminal DNA binding domain (DBD) for binding auxin response elements (AuxREs) and ARF dimerization, a middle region (MR) and the PB1 domain. ARFs can be divided into two functional groups based on their amino acid composition in the middle region (MR) and their activities as transcription activators or repressors in transient expression assays. The MRs of class A ARFs (also called activator ARFs) have a high glutamine content while classes B and C ARFs (also called repressor ARFs) are rich in serine, threonine and proline residues^18,19^. The canonical AFB/IAA/ARF signaling cascade described above is mainly based on class A ARFs whose members include ARF5, ARF6, ARF7, ARF8 and ARF19 in *Arabidopsis thaliana*. Recent studies in moss and liverwort suggest that class B ARFs may function as transcription repressors by competing with class A ARFs for binding to auxin-responsive promoters^20,21^. Class C ARFs are also transcription repressors, although they may regulate auxin-independent development^21^.

The mechanism of auxin-induced fruit set has been studied by genetic analyses. In multiple species, mutations or silencing of class A ARF(s) leads to parthenocarpic fruits from emasculated flowers, supporting the repressive role of class A ARFs in ovary-derived fruit set^1,7,22–25^. However, by characterizing higher order mutant combinations of four class A *SlARF*s (*SlARF5, SlARF7, SlARF8A, SlARF8B*) in tomato (*Solanum lycopersicum*), we recently demonstrated that these class A SlARFs display dual functions during fruit development^26^. Before pollination, all four SlARFs act as inhibitors of fruit set when associated with SlIAA9. After fertilization, the elevated auxin levels in the ovary result in SlIAA9 degradation and free the class A SlARFs to function as activators in subsequent fruit growth. The positive role of SlARFs in fruit growth is reflected by the biphasic bell-shape curve of parthenocarpic fruit size in response to varying doses of these SlARFs. The maximum parthenocarpic fruit size was reached by removal of SlARF8A and SlARF8B, while knocking out all four SlARFs abolished fruit growth completely. Consistent with this idea, expression of truncated SlARF8A or 8B lacking the SlIAA9-interacting PB1 domain in transgenic tomato resulted in the production of large seedless fruits^26^. Similarly, AtARF6 and AtARF8 in *Arabidopsis* also showed dual role in fruit initiation/growth, although the specific IAA(s) regulating this process have not been identified^27^. The *arf6* and *arf8* single mutants produced longer and wider fruits from emasculated unfertilized flowers^22,27^. In contrast, the *arf6 arf8* double mutant produced short pistils with similar length as that of WT without fertilization^27^. These observations suggest that although both ARF6 and ARF8 inhibit fruit set before anthesis, they then promote fruit elongation after fertilization.

Class A ARFs in *Arabidopsis* have been shown to be dynamically regulated by post-translational modifications (PTMs), including phosphorylation, Small Ubiquitin-Like Modifier (SUMO)-modification (SUMOylation), and ubiquitination^18,28^. Phosphorylation of ARF7/19 MRs, which is induced by a peptide hormone TRACHEARY ELEMENT DIFFERENTIATION INHIBITORY FACTOR (TDIF), enhances ARF DNA binding affinity and reduces interaction with IAA proteins to promote lateral root formation^29^. SUMOylation of ARF7 contributes to hydropatterning of the *Arabidopsis* root in response to moisture by promoting ARF7-IAA3 interaction to inhibit lateral root initiation specifically in dry environments^30^. ARF6 ubiquitination promotes its degradation in response to abscisic acid (ABA) treatment in *Arabidopsis* seedlings^31^. However, whether and how PTMs regulate ARF6- and ARF8-mediated fruit set/growth were unknown. In this study, we reveal that both ARF6 and ARF8 are *O*-fucosylated by SPINDLY (SPY), a novel nucleocytoplasmic protein *O-*fucosyltransferase (POFUT), which catalyzes the addition of a fucose moiety to the hydroxyl oxygen of specific Ser/Thr residues of target proteins^32^. Importantly, *O*-fucosylation of MRs of ARF6/8 reduced their binding to MED8, a subunit of the coactivator Mediator complex. The pleiotropic phenotypes of the *spy* mutants and recent discovery of hundreds of SPY target proteins by proteomic studies all point to the important roles of SPY in regulating diverse cellular processes^33–35^. But the molecular mechanism of SPY regulation has only been defined for a handful of its targets^32,34,36–39^. Here, we demonstrated the role of SPY in attenuating auxin-induced fruit set and growth by *O*-fucosylating ARF6 and ARF8, and their interacting proteins, co-repressor IAA9 and MED8 of the coactivator Mediator complex.

## RESULTS

### *spy* mutants displayed parthenocarpic fruit growth

Genetic and proteomic analyses indicate that protein *O*-fucosylation by the nucleocytoplasmic-localized SPY regulates diverse developmental processes in *Arabidopsis*, although most of the molecular mechanisms are unknown^33–35,40^. Two previous mutant studies investigated the effect of *spy* mutations on parthenocarpy with contradicting results. One study reported that *spy-2* and *spy-3* in the Col-0 ecotype displayed parthenocarpy after emasculation ^41^, while another study did not observe any parthenocarpy phenotype in *spy-3* (Col-0 ecotype) or *spy-4* (Ws ecotype)^42^. To clarify whether SPY plays a role in parthenocarpy, we examined the pistil phenotypes of four *spy* alleles, including *spy-8* and *spy-19* in the L*er* background and *spy-3* and s*py-23* in the Col-0 background. All four *spy* mutants showed significantly longer pistils from emasculated flowers comparing to those of wild-type controls (**Fig. 1a-1d**). Furthermore, the pistil phenotype of *spy-3* was partially rescued by introduction of *P_SPY_:GFP-SPY* or *P_SPY_:GFP-SPY-NLS* (nuclear localization sequence), but not by *P_SPY_:GFP-SPY-NES* (nuclear export sequence) (**Fig. 1e-1f**, **Supplementary Fig. 1a-1b**). These results indicate that nuclear-localized SPY inhibits fruit initiation and elongation before pollination. Consistent with this notion, we found that SPY protein levels were reduced after anthesis as detected using a *P_SPY_:FLAG-SPY spy-3* transgenic line (**Supplementary Fig. 1c**). We also examined overall protein *O*-fucosylation before and after anthesis by protein blot analysis using a biotinylated *Aleuria aurantia* lectin (AAL), a terminal fucose-specific lectin. The *P_SPY_:FLAG-SPY spy-3* line showed reduced protein *O*-fucosylation at 3 DAA and 5 DAA compared to that at –2 DAA and 0 DAA (**Supplementary Fig. 1d**).

**Figure 1.**
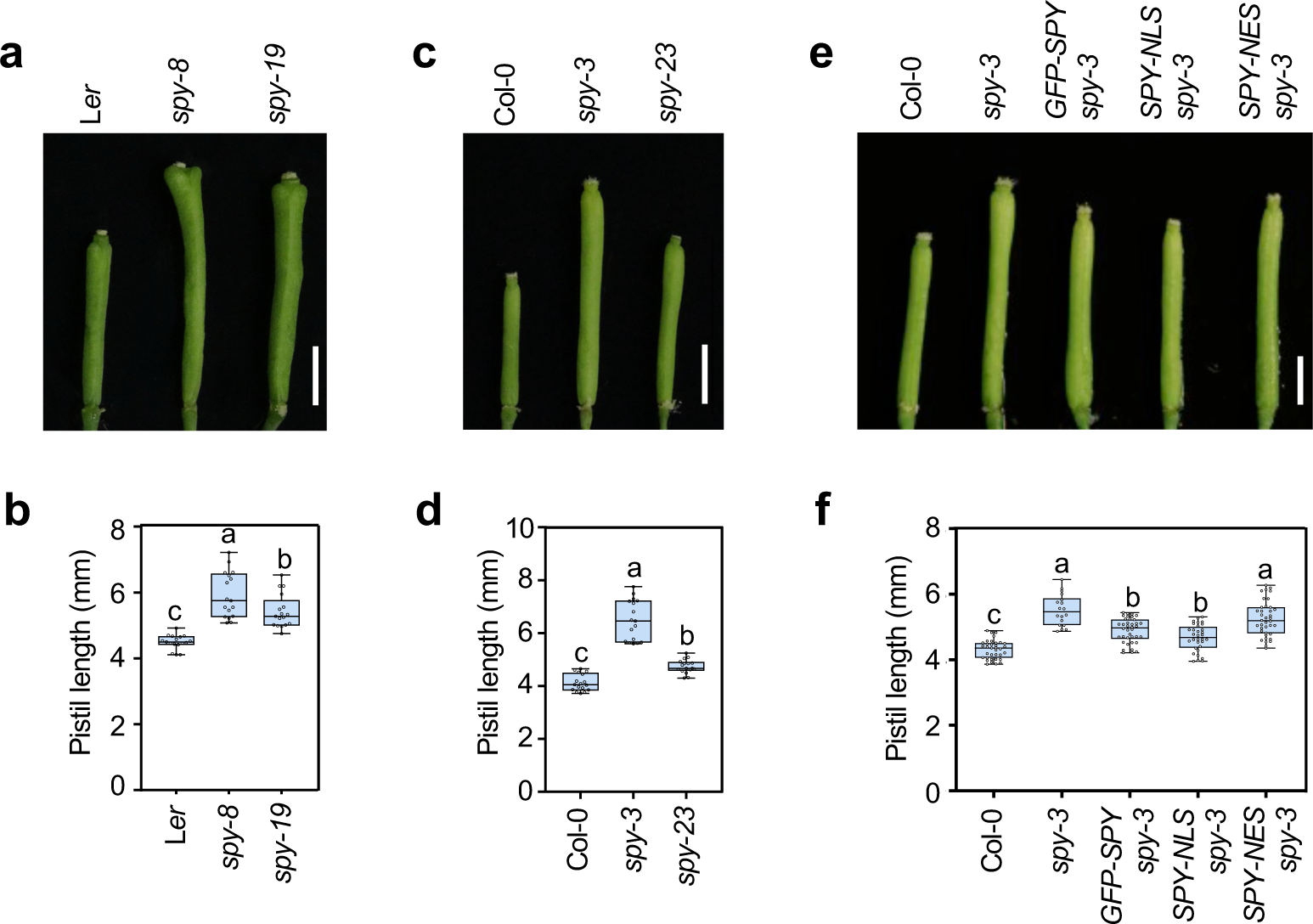
*spy* mutations promoted parthenocarpic fruit growth. **a-d**, *spy* mutations in both L*er* and Col-0 backgrounds promoted parthenocarpic fruit growth. In **a** and **c**, photo showing representative pistils of different genotypes 7 days after emasculation. Bar = 2 mm. In **b** and **d**, pistil lengths. n>15. **e-f**, the nuclear-localized SPY partially rescued the pistil phenotype of *spy-3*. In e, Bar = 2 mm. In **f**, pistil lengths. n>15. GFP-SPY, GFP-SPY-NLS and GFP-SPY-NES were accumulated at similar levels in these lines (**Supplementary Fig. 1b**). In boxplots **b**, **d** and **f**, center lines and box edges are medians and the lower/upper quartiles, respectively. Whiskers extend to the lowest and highest data points within 1.5x interquartile range (IQR) below and above the lower and upper quartiles, respectively. Different letters above the boxes represent significant differences (*p* < 0.05) as determined Tukey’s honestly significant difference (HSD) mean separation test. Two biological repeats showed similar results.

### *arf6* showed additive interaction with *spy*, whereas *arf8* was epistatic to *spy* in parthenocarpic growth

The *arf6* and *arf8* mutants were shown previously to display longer and wider fruit after emasculation comparing to that of WT ^27^. Consistent with the previous report, we found that both *arf6-2* and *arf8-3* showed increased fruit length and width after emasculation, although *arf8* displayed stronger parthenocarpic growth than *arf6* (**Fig. 2a, 2b, and Supplementary Fig. 2a**). These pistil phenotypes were partially rescued by *P_ARF6/8_:FLAG-ARF6/8* (**Supplementary Fig. 3a, 3b**). Another class A ARF, ARF7 (also known as NPH4), is also expressed in the developing pistil (Arabidopsis eFP Browser^43^, http://bar.utoronto.ca/efp_arabidopsis/cgi-bin/efpWeb.cgi). However, the *nph4-1* mutation alone or in combination with *arf6* and/or *arf8* did not increase fruit growth after emasculation (**Supplementary Fig. 2b, 2c**), suggesting that *ARF7* is not a major regulate of this process.

**Figure 2.**
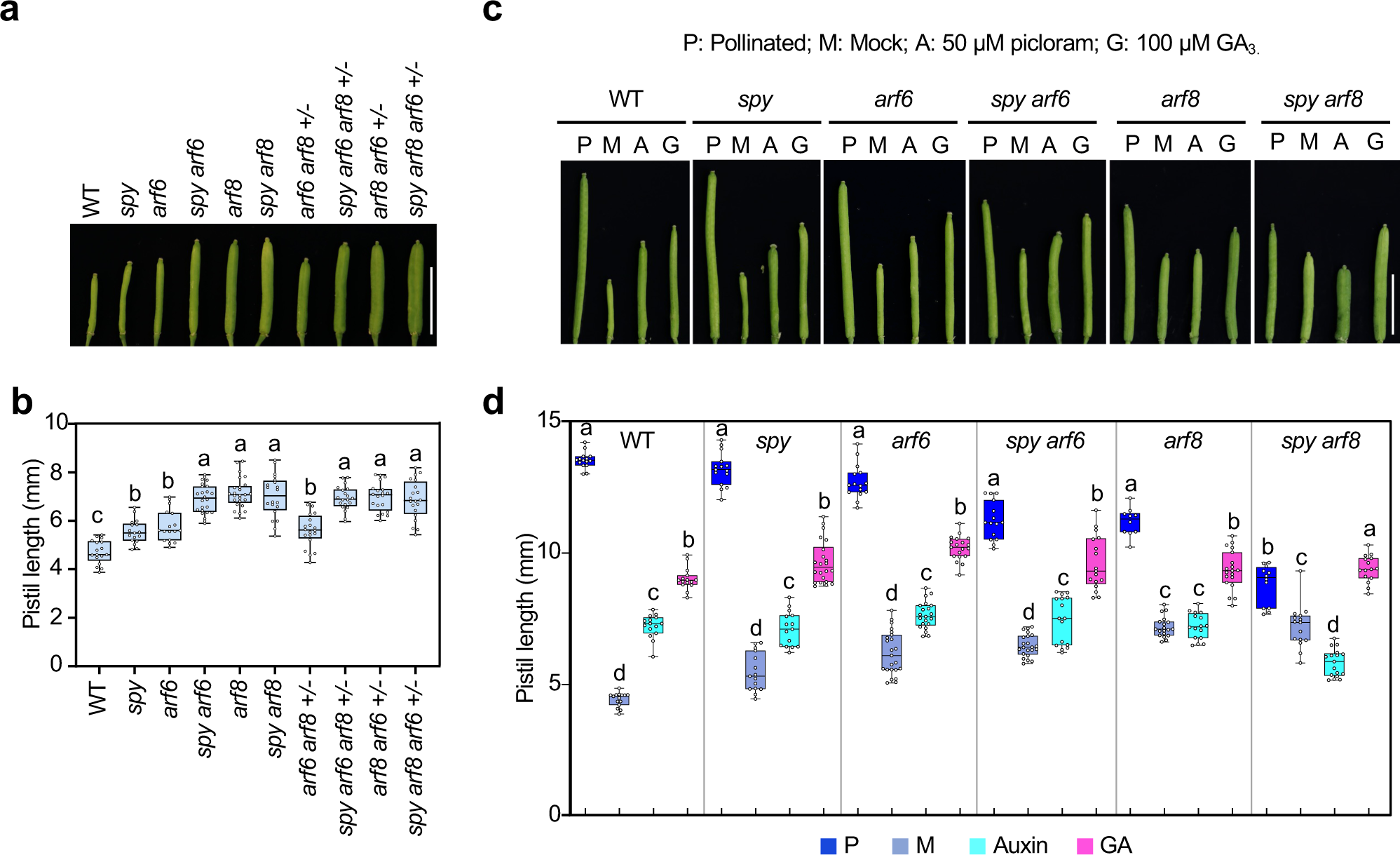
ARF6 and ARF8 mediated auxin-induced parthenocarpic growth downstream of SPY. **a-b**, epistasis analysis of *spy-3*, *arf6-2* and *arf8-3* mutations. *spy-3* and *arf6* additively promoted parthenocarpic fruit elongation, whereas *spy-3 arf8* showed similar phenotype as *arf8*. Mutant alleles are homozygous unless specified as heterozygous, including *arf6* +/- and *arf8* +/-. In **a**, photo showing representative pistils of different genotypes 7 days after emasculation. Bar = 5 mm. In **b**, pistil lengths. n>15. **c-d**, the *spy arf6* double mutant was responsive to picloram treatment, whereas *spy arf8* was not. All mutants were responsive to GA. In **c**, photo showing representative pistils of different genotypes 7 days after emasculation and treatment with mock solvent (M) or 50 µM picloram (A, for auxin analog) or 100 µM GA_3_ (G). Pistils from self-pollinated flowers (P) were included for comparison. Bar = 5 mm. In **d**, pistil lengths. n>15. In boxplots **b** and **d**, center lines and box edges are medians and the lower/upper quartiles, respectively. Whiskers extend to the lowest and highest data points within 1.5x IQR below and above the lower and upper quartiles, respectively. Different letters above the boxes represent significant differences (*p* < 0.05) as determined by Tukey’s HSD mean separation test. Two biological repeats showed similar results.

To examine the genetic interaction between *SPY* and *ARF6*/*8*, epistasis analysis was performed among *spy-3*, *arf6-2* and *arf8-3* using single, double, triple mutants. Most of the mutant alleles are homozygous, except that mutants containing both *arf6* and *arf8* mutations were sesquimutants (*arf6 -/- arf8 +/-* and *arf8 -/- arf6 +/-*) because the *arf6 arf8* double homozygote is sterile. The *spy*, *arf6* and *arf8* single homozygous mutants produced longer pistils than that of WT with *arf8* displayed the strongest phenotype (**Fig. 2a-2b**). Importantly, we found that *spy* and *arf6* additively promoted parthenocarpic fruit elongation in the *spy arf6* double homozygous mutant. Similarly, in the *arf6 arf8+/-* sesquimutant background, the *spy* mutation further increased fruit length. However, the homozygous double mutant *spy arf8* showed similar fruit length as that of *arf8*. In the *arf8 arf6+/-* sesquimutant background, the *spy* mutation did not further enhance fruit growth either. These results indicated that *arf8* is epistatic to *spy*, suggesting that ARF8 is downstream of SPY in regulating parthenocarpy. The additive interaction between *spy* and *arf6* in parthenocarpic growth is likely because both ARF6 and ARF8 are downstream of SPY, but ARF8 plays a more predominant role in this process. This notion is further supported by our results in the next section showing that both ARF6 and ARF8 are *O*-fucosylated by SPY.

Besides auxin, previous studies show that another phytohormone gibberellin (GA) also promotes fruit initiation and growth in *Arabidopsis*. The quadruple or global *della* mutant (knockout four or all 5 *DELLA* genes) displays strong parthenocarpy^6,44^. Because SPY represses GA responses in hypocotyl elongation by enhancing DELLA activity via *O*-fucosylation, we tested whether SPY’s regulation of parthenocarpy is through its repression of both GA and auxin responses by comparing GA vs auxin responses in parthenocarpic fruits of WT and different mutant backgrounds. Application of GA_3_ or the auxin analog picloram^45^ to pistils of emasculated WT flowers promoted parthenocarpic fruit growth (**Fig. 2c-2d**). Treatment of GA or auxin also increased parthenocarpic fruit growth in *spy*, *arf6*, and *spy arf6*. However, only GA treatment, but not auxin, increased fruit growth in *arf8* or *spy arf8*. These results indicate that SPY represses parthenocarpy by regulating both GA and auxin responses, and that ARF8 plays a more predominant role than ARF6 in mediating auxin-induced parthenocarpic growth downstream of SPY.

### ARF6 and ARF8 are *O*-fucosylated by SPY

Based on the genetic interaction between *spy* and *arf6/8*, we tested whether ARF6 and ARF8 are direct targets of SPY. By transient co-expression of FLAG-ARF6/ARF8 with SPY in *Nicotiana benthamiana*, we found that both affinity-purified FLAG-ARF6 and -ARF8 were *O*-fucosylated as detected by biotinylated fucose-specific lectin, AAL (**Supplementary Fig. 4a-4b**). To confirm that ARF6 and ARF8 are *O*-fucosylated in *Arabidopsis*, we generated transgenic *Arabidopsis* lines carrying either *P_UBQ10_:FLAG-ARF6* or *P_UBQ10_:FLAG-ARF8*, and then introduced these transgenes into the *spy-8* background. Affinity purified FLAG-ARF6 and - ARF8 proteins from the WT background, but not those from the *spy-8* background, were *O*-fucosylated as detected by AAL-biotin (**Fig. 3a-3b**). To identify the *O*-fucosylation (Fuc) sites in these proteins, affinity-purified FLAG-ARF6 and -ARF8 from *N. benthamiana* and transgenic *Arabidopsis* were analyzed by liquid chromatography (LC)-electrospray ionization (ESI)-mass spectrometry (MS). One *O*-fucosylated ARF6 peptide and three *O*-fucosylated ARF8 peptides were identified (**Fig. 3c-3d**, **Supplementary Table 1, Supplementary Data Sets 1-4**). Notably, all identified *O*-Fuc sites are in the MR of ARF6/8. We also performed an AAL pulldown assay using *N. benthamiana* that transiently expressed full-length or truncated ARF6 proteins in the presence or absence of SPY. Besides the full-length ARF6, only the MR fragment (amino acid residues 376-778) but not the N-terminal DBD-containing fragment (amino acid residues 1-375) or C-terminal PB1 domain (amino acid residues 779-935) was *O*-fucosylated (**Fig. 3e**). These results support that the major *O*-Fuc sites in ARF6 and ARF8 reside within their MR sequence.

**Figure 3.**
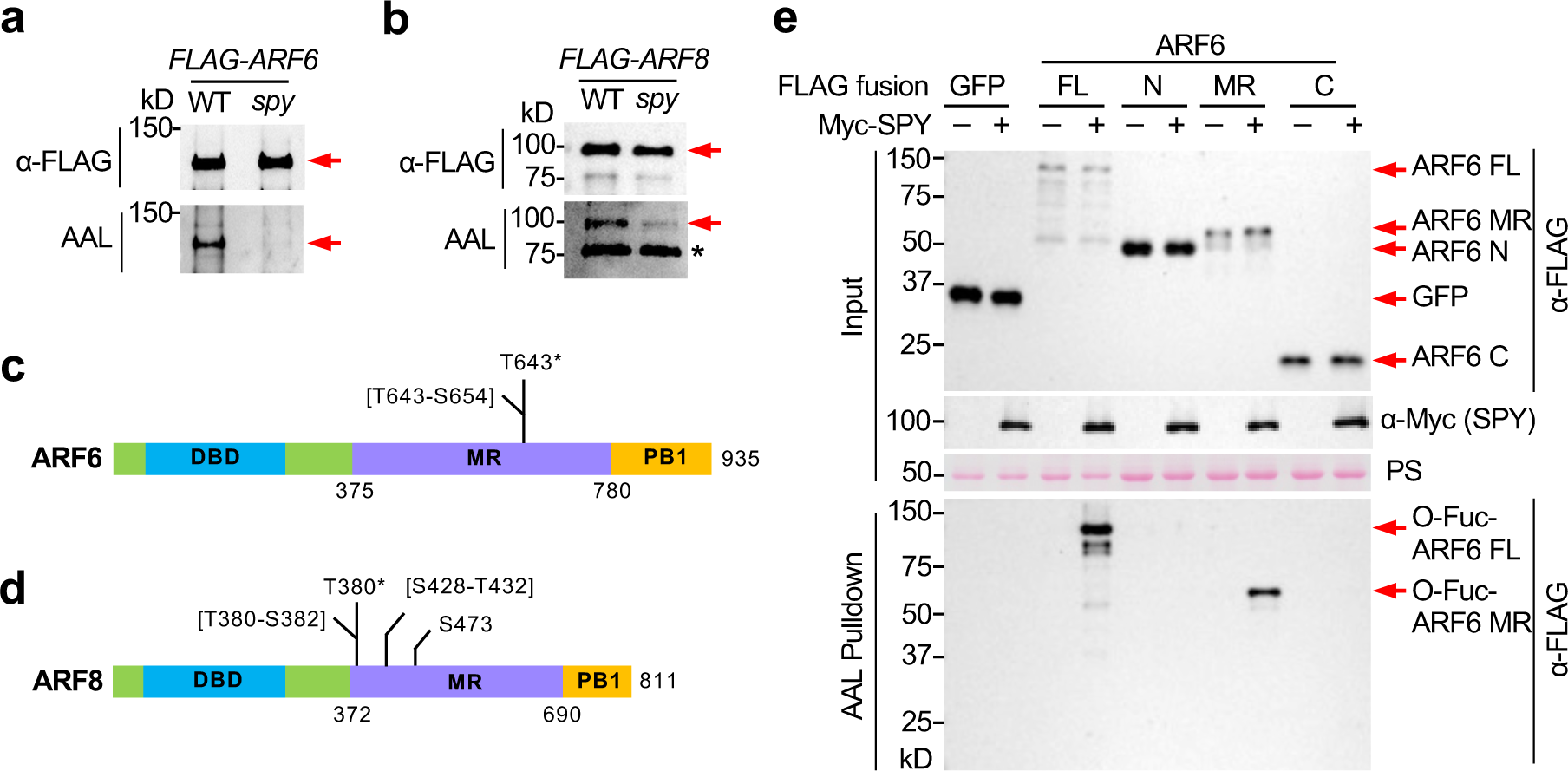
Identification of *O*-fucosylation sites in ARF6 and ARF8. **a**, FLAG-ARF6 was *O*-fucosylated by SPY in *Arabidopsis*. **b**, FLAG-ARF8 was *O*-fucosylated by SPY in *Arabidopsis*. In **a**-**b**, FLAG-ARF6/8 proteins were affinity-purified from transgenic *Arabidopsis* carrying *P_UBQ10_:FLAG-ARF6* or -*ARF8* in WT or *spy-8* background, and protein blots were probed with either AAL-biotin or anti-FLAG antibody as labeled. In **a-b**, arrow in the top panel indicates FLAG-ARF6 or -ARF8, and arrow in the bottom panel indicates *O*-fucosylated FLAG-ARF6 or -ARF8. In **b**, * indicates a non-specific background band. **c-d,** ARF6 and ARF8 *O*-fucosylation sites identified by MS analysis. The schematic shows the ARF6 (**c**) or ARF8 protein (**d**); The marked S/T residues are confirmed *O*-Fuc sites. The sequence within square brackets contains undetermined *O*-Fuc sites. *, also identified in a recent proteomic study^35^. **e,** AAL pulldown assay confirmed that MR-ARF6 contains major *O*-Fuc site(s). FLAG-tagged full-length (FL) or truncated ARF6 proteins were expressed alone (–) or co-expressed (+) with Myc-SPY in *N. benthamiana.* FLAG-GFP, a negative control. *O*-fucosylated proteins were pull-downed by AAL-agarose. Immunoblot containing input (top panel) or AAL-agarose pull-down samples (bottom panel) was probed with anti-FLAG and anti-Myc antibodies as labeled. PS, Ponceau S-stained blot showing even loading. N, N-terminal DBD domain; MR, middle region; C, C-terminal PB1 domain of ARF6. Two biological repeats showed similar results.

In addition to *O*-fucosylation, we found that both ARF6 and ARF8 contained two other types of PTMs, phosphorylation and *O*-link-*N*-acetylglucosamine (GlcNAc) modification (**Supplementary Table 1 and Supplementary Fig. 5**). Intracellular protein *O*-GlcNAcylation is catalyzed by an *O*-GlcNAc transferase (OGT), SECRET AGENT (SEC) that is a paralog of SPY in *Arabidopsis*^46^. Recent proteomic studies showed that SPY and SEC have unique targets as well as common targets in *Arabidopsis*^34,35,47,48^. The interplay between SPY and SEC is complex as these two types of glycosylation can interact antagonistically or additively, depending on the target proteins^32,38,46,49^. We found that the *sec* mutants produced short pistils after emasculation with similar length as that of WT (**Supplementary Fig. 4c, 4d**), suggesting that *O*-GlcNAcylation does not alter the function of ARF6/8 during fruit set significantly. Therefore, we focused on the regulatory mechanism of SPY on ARF6/ARF8 in the rest of this study.

### SPY and ARF6/8 regulate many common target genes during fruit set

Our genetic and biochemical analyses indicated that ARF6 and ARF8 are downstream of SPY in regulating parthenocarpy. To identify ARF6-, ARF8- and SPY-responsive genes that are involved in fruit set and growth, transcriptome analysis was performed by RNA-seq using the following samples: (1) unpollinated pistils at 2 days before anthesis (−2 DAA, from stage 10 flowers) of *arf6-2, arf8-3, spy-3* and WT (Col-0); and (2) 0 DAA WT pistils (from stage 14 flowers) that were already self-pollinated. Two biological repeats were included in each set of samples. The differentially expressed gene (DEG) lists for ARF6, ARF8 or SPY-responsive genes (108, 1143, 271 DEGs, respectively) were identified by comparing each mutant vs WT (– 2 DAA) dataset using the criteria of fold change > 1.5 and *p* < 0.05 (**Fig. 4a**, **Supplementary Fig. 6a, Supplementary Table 2**). The DEG list for fertilization-responsive genes (2528 total) was generated by comparing 0 DAA WT vs –2 DAA WT dataset with fold change > 1.5 and *p* < 0.05 (**Supplementary Fig. 6a, Supplementary Table 2**). Consistent with the stronger parthenocarpic phenotype of *arf8*, the *arf*8 mutation resulted in altered transcript levels of many more genes than *arf6*, and almost all ARF6 DEGs (99 out of 108 total) were included in ARF8 DEG list (**Fig. 4a**). Comparison of DEG lists for SPY, ARF6 and ARF8 revealed that 67% of SPY DEGs (181 DEGs out of 271 total) were co-regulated by ARF6 and/or ARF8 (**Fig. 4a**, **Supplementary Table 3**). Among these co-regulated DEGs, 67 genes were up-regulated by both *spy-3* and *arf8* (**Fig. 4b, 4d**, **Supplementary Table 3**), and 103 genes were down-regulated by both *spy-3* and *arf8* (**Fig. 4c, 4d**, **Supplementary Table 3**). Comparison of SPY- and ARF8-responsive genes with fertilization-responsive genes (0 DAA WT vs –2 DAA WT) further showed that 62% of SPY- and ARF-coregulated genes were also fertilization-responsive genes (113 DEGs out of 181 total, **Supplementary Fig. 6b-6e**). The significant overlap among the SPY-, ARF6/ARF8-, and fertilization-responsive genes provide strong support that SPY regulates fruit set at least in part mediated by ARF6 and ARF8.

**Figure 4.**
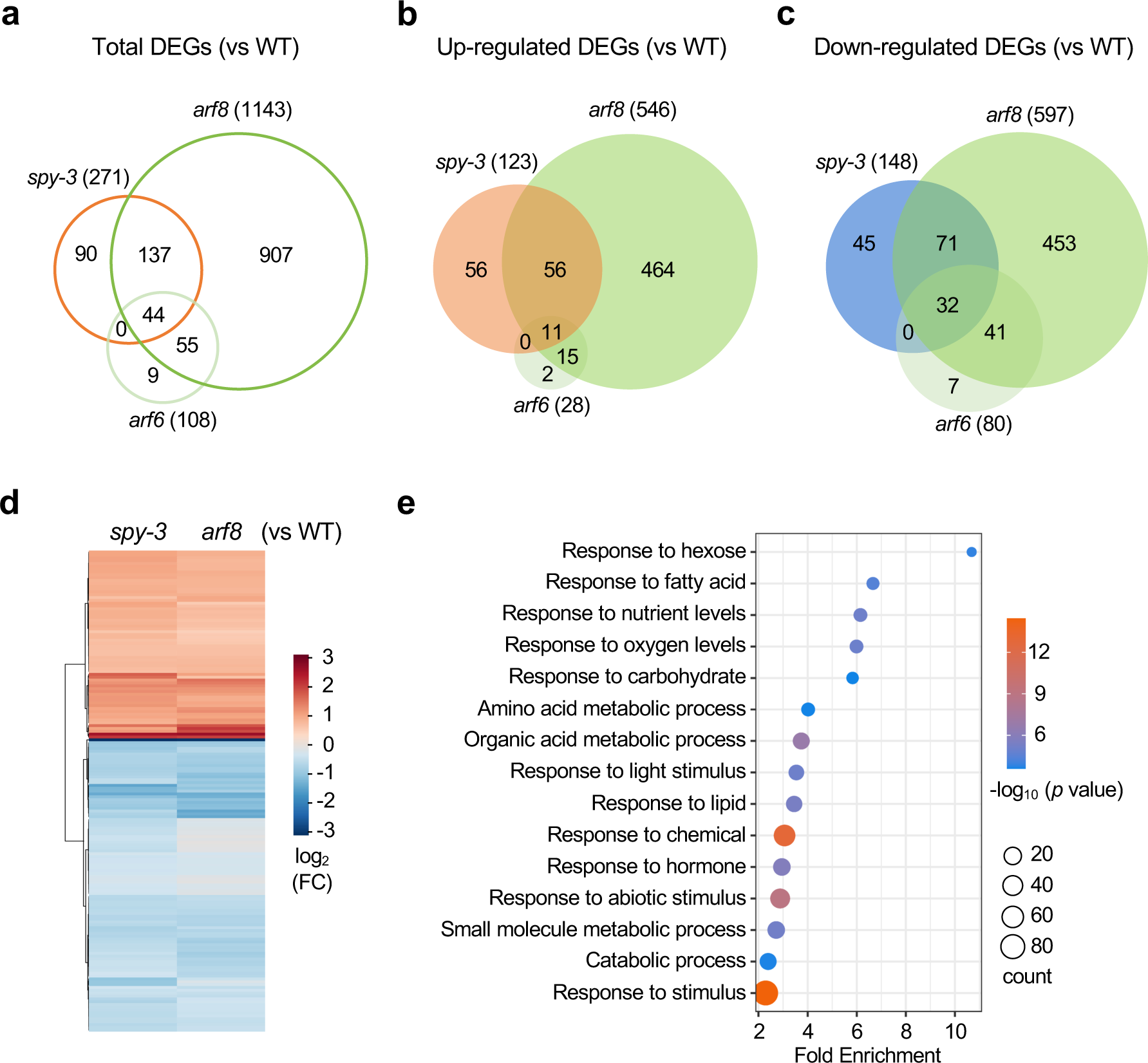
Identification of ARF6-, ARF8- and SPY-responsive genes in pistils by RNA-seq analysis. RNA-seq analysis was performed using −2 DAA pistils (stage 10) of *arf6, arf8, spy-3* and WT. The differentially expressed gene (DEG) lists for ARF6-, ARF8- and SPY-responsive genes are in **Supplementary Table 2**. **a-c,** Venn diagrams of coregulated DEGs by *ARF6, ARF8* and *SPY*. Total DEGs in **a**, Up-regulated DEGs in **b**, Down-regulated DEGs in **c**. **d**, Heat map of SPY and ARF8 coregulated 181 DEGs. **e**, Enrichment of selected biological processes in ARF8 and SPY co-regulated 181 DEGs by GO term analysis.

Gene Ontology (GO) analysis of SPY- and ARF6/8-coregulated DEGs identified 48 enriched GO terms (**Supplementary Table 4**). Although the coregulated DEGs belong to a variety of biological processes, certain groups were more represented, including response to sugar, fatty acid, nutrient and hormone, and biogenesis related genes for carbohydrate/amino acid metabolic processes (**Fig. 4e**). Seven SPY- and ARF-coregulated genes were selected to verify their expression in –2 DAA WT, *spy-3* and *arf8*, and 0 DAA WT by RT–qPCR (**Supplementary Table 5**). Among them, three genes encode transcription repressor or transcription factor: *IAA19* in auxin signaling^50,51^, *ETHYLENE RESPONSE FACTORs* (*ERF107* and *ERF023*) that are responsive to oxylipins^52^. In addition, ERF107 regulates nitrate assimilation^53^, and *ERF023* is responsive to cellular nitrogen status^54^. Two genes are involved in sugar response/metabolic process, and their expression is repressed by sugar: *BASIC LEUCINE-ZIPPER 1* (*bZIP1*) encoding a S1 subgroup bZIP transcription factor^55,56^ and *GNTL* encoding a beta-1,6-N-acetylglucosaminyl transferase-like enzyme for the synthesis of glycan and/or glycosylation of proteins^57–59^. *ARABINOGALACTAN PROTEIN 12* (*AGP12*) was implicated in nutrient uptake in developing seeds^60^, and *SHORT LIFE* (*SHL)* encodes a plant-homeodomain (PHD) protein that functions as a histone reader for chromatin remodeling in repressing floral induction and regulating fertility^61–63^. The RT–qPCR assays confirmed that *AGP12*, *ERF023* and *IAA19* were upregulated, whereas *GNTL*, *ERF107*, *bZIP1* and *SHL* were downregulated in *spy-3*, *arf8* or by fertilization (WT 0 DAA) in comparison to that in WT –2 DAA (**Fig. 5a, 5b**). Up-regulation of *IAA19* expression by *arf8* and *spy* reflects elevated auxin response in these mutant pistils. AGPs are known to function in cell wall reorganization^57^. Induction of *AGP12* by *arf8* and *spy* is consistent with the proposed role of this gene in nutrient uptake of ovule based on its expression at the chalaza of the ovule and funiculus^60^. Auxin has been shown to promote sugar transport and metabolism in ovaries after fertilization in several species^64–66^, which in turn inhibits fruit abortion caused by programmed cell death and promotes fruit growth^64^. In *Arabidopsis* pistils at –2 DAA, we found that two sugar-repressed genes *bZIP1* and *GNTL* were downregulated by *arf8* and *spy*, suggesting that these two mutations led to an increased sugar content/signaling and that bZIP1 and GNTL may play a negative role in fruit set/growth in WT before anthesis. Consistent with this idea, two *bZIP1* knockout mutants displayed longer pistils after emasculation comparing to WT (**Supplementary Fig. 7a, 7b**).

**Figure 5.**
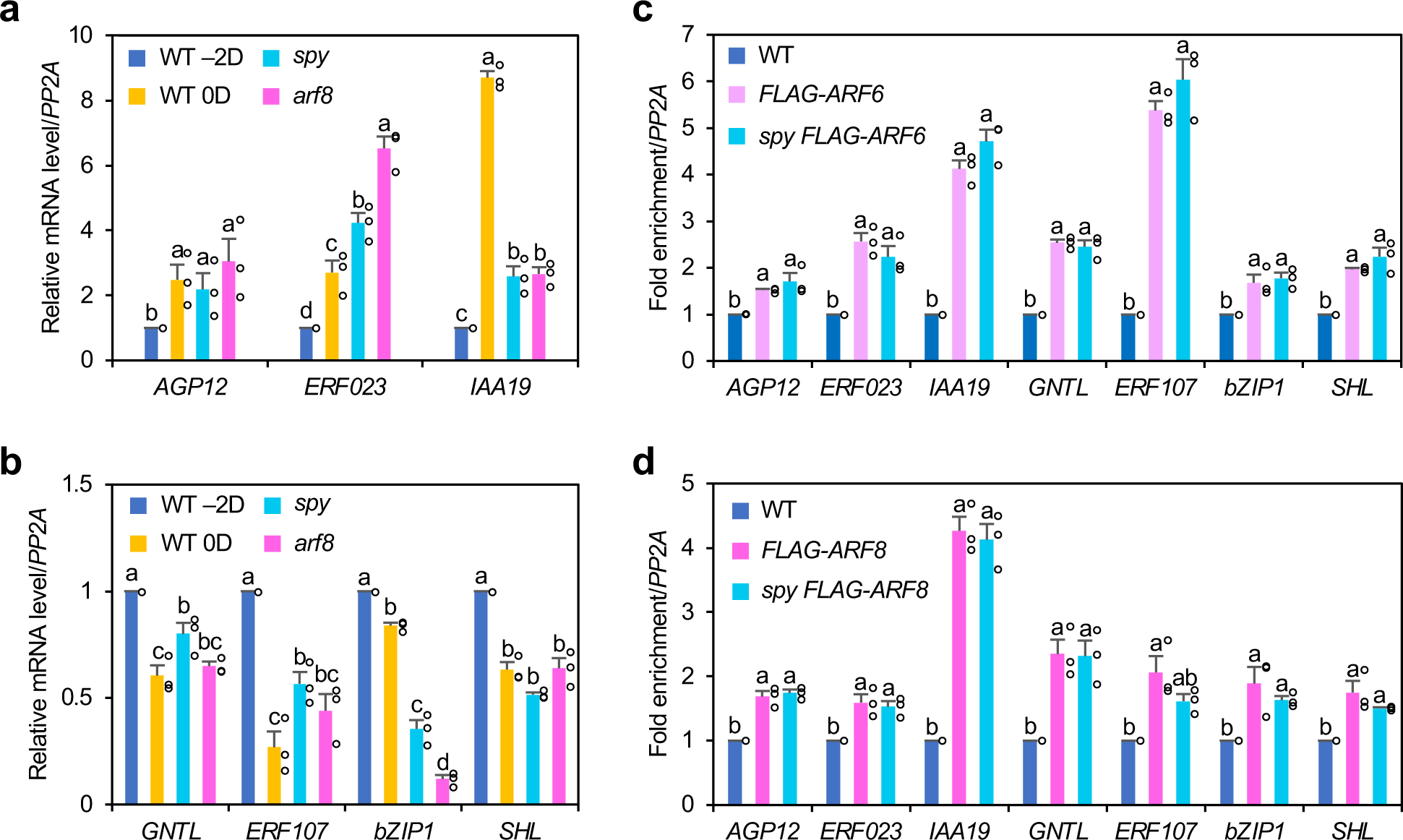
Confirmation of *ARF6/ARF8* target genes by RT-qPCR and ChIP-qPCR. **a-b**, RT-qPCR analysis confirming selected genes that were upregulated (in **a**) or downregulated (in **b**) in −2 DAA pistils of *arf8* and *spy-3* mutants in comparison to WT. These genes were also upregulated or downregulated, respectively, after pollination (0 DAA WT vs –2 DAA WT pistils). For all RT-qPCR analyses, the housekeeping gene *PP2A* was used to normalize different samples. Means ± SE of 3 biological replicas are shown. Expression level in –2 DAA WT pistil was set to 1. **c-d**, ChIP-qPCR analysis showed ARF6 (in **c**) and ARF8 (in **d**) binding to promoter regions of selected ARF-responsive genes, although *spy* mutation did not affect ARF binding. −2 DAA pistils of the *P_UBQ10_:FLAG-ARF6/ARF8* lines in WT or *spy* mutant backgrounds and anti-FLAG beads were used for the ChIP experiment. The relative enrichment was calculated by normalizing against ChIP-qPCR of non-transgenic WT samples using *PP2A* as control. Means ± SE of 3 biological replicas are shown. In **a-d**, Different letters above the bars represent significant differences (*p* < 0.05) as determined by Tukey’s HSD mean separation test.

To identify direct target genes of ARF6/ARF8 that are coregulated by SPY, we compared our SPY- and ARF6/ARF8-responsive DEG list (181 total) with a published ARF6 ChIP-seq dataset generated using seedling samples^51^. This comparison identified 51 overlapping genes as direct targets of ARF6/ARF8, including six of the selected seven genes (except *bZIP1*) verified by RT-qPCR (**Supplementary Table 6**, **Supplementary Fig. 6f**). Although bZIP1 is not present in the published ARF6 ChIP-seq dataset, its promoter region contains tandem AuxREs. It is possible that bZIP1 is a direct ARF target in the pistil, but not at the seedling stage. ChIP-qPCR assay was performed to verify direct binding of ARF6/8 to the promoters of these genes. Transgenic *Arabidopsis* seedlings carrying either *P_UBQ10_:FLAG-ARF6* or *P_UBQ10_:FLAG-ARF8* in the WT background were used for chromatin crosslinking and immunoprecipitation using anti-FLAG beads. The non-transgenic WT was included as the negative control. qPCR was performed using primers that span the ARF6 binding peaks in the promoter of each target gene, except for *bZIP1* where primers span two tandem AuxREs in its promoter. The ChIP-qPCR analysis showed a significant enrichment in all seven target promoters in the *P_UBQ10_:FLAG-ARF6* and *P_UBQ10_:FLAG-ARF8* lines compared to the WT control (**Fig. 5c, 5d**), supporting that these genes are direct targets of ARF6 and ARF8.

### SPY did not affect ARF6 or ARF8 DNA binding, protein stability or nuclear localization

Because SPY and ARF6/ARF8 coregulate many target genes during fruit set, we hypothesized that *O*-fucosylation of ARF6 and ARF8 by SPY may enhance ARF6/8 function to inhibit fruit set. To investigate the molecular mechanism involved, we first examined whether this posttranslational modification increases binding of ARF6/ARF8 to their target promoters by ChIP-qPCR using transgenic *P_UBQ10_:FLAG-ARF6* or *P_UBQ10_:FLAG-ARF8* lines in WT or *spy* background. We did not observe significant reduction in enrichment of target promoter sequences by the *spy* mutation (**Fig. 5c, 5d**), suggesting that SPY does not alter the DNA binding activities of ARF6/8. Recent studies have reported that the protein stability and nucleocytoplasmic partitioning of two class A ARFs, ARF7 and ARF19, in *Arabidopsis* are differentially regulated during root development^67,68^. By immunoblot analysis and confocal fluorescence microscopy, we showed that FLAG-or GFP-tagged ARF6 and ARF8 proteins accumulated to similar levels in the WT and *spy* backgrounds (**Supplementary Fig. 8a-8d**). Unlike the nucleocytoplasmic partitioning reported for ARF7 and ARF19^67^, ARF6 was only detected in the nucleus in roots and pistils.

### SPY enhanced IAA9-ARF6/8 transcription repression, but reduced ARF6/8 transactivation activity

As described in the Introduction, class A ARFs play dual function in fruit development in tomato and in *Arabidopsis*. In tomato, four class A-SlARFs together with SlIAA9 inhibit fruit set but promote subsequent fruit growth after fertilization when elevated auxin levels trigger SlIAA9 degradation ^26^. ARF6 and ARF8 in *Arabidopsis* also showed similar dual function in fruit development ^27^. We reasoned that *AtIAA9* is likely the functional ortholog in this process because its mRNA levels are elevated in ovaries (pistils) before anthesis^69^ (Arabidopsis eFP Browser^43^, http://bar.utoronto.ca/efp_arabidopsis/cgi-bin/efpWeb.cgi) and IAA9 has been reported to interact with ARF6 and ARF8 to repress auxin-induced adventitious root initiation^70^. Consistent with this idea, the *iaa9-1* mutant produced slightly longer pistils after emasculation compared to WT (**Supplementary 7c, 7d**). The subtle parthenocarpic phenotype of *iaa9-1* is likely because the T-DNA insertion site in this allele is located in its fourth intron^71^, which is predicted to cause C-terminal truncation at the end of the PB1 domain. To examine the effects of IAA9 and SPY on the transcription activities of ARF6 and ARF8, dual luciferase (LUC) assays^72^ were performed using the transient expression system in *N. benthamiana*. A synthetic auxin-responsive promoter *P3(2x)* was fused to the *firefly LUC (fLUC)* as the reporter for this assay because *P3(2x)* was shown to be responsive to AtARFs^73^. *35S:Renilla LUC (rLUC)* was used as the internal control to normalize variations in transformation efficiency. Five effectors, *35S:FLAG-ARF6*, *35S:FLAG-ARF8, 35S:Myc-IAA9*, *35S:HA-SPY* and *35S:HA-(spy-19)* were included in the assays. A catalytic-domain mutant allele, *spy-19*^32^, served as a negative control. To avoid variations in the IAA9 protein levels, the *IAA9* construct used in this assay encodes a dominant stabilized mutant protein with a P188S substitution in the conserved degron to prevent auxin-induced degradation^74–76^. As expected, expression of ARF6 or ARF8 alone induced *P3(2x):fLUC* transcription. Importantly, co-expression of SPY, but not spy-19, attenuated transactivation activities of both ARFs (**Fig. 6a, 6b,** and **Supplementary Fig. 9**). Co-expression of ARF6 or ARF8 with IAA9 repressed *P3(2x):fLUC.* Notably, co-expression of ARF, IAA9 and SPY further repressed *P3(2x):fLUC.* These results suggested that *O*-fucosylation by SPY reduced ARF6/8 transactivation activity, while enhanced the transcription repression activity of the IAA9-ARF6/8 complexes.

**Figure 6.**
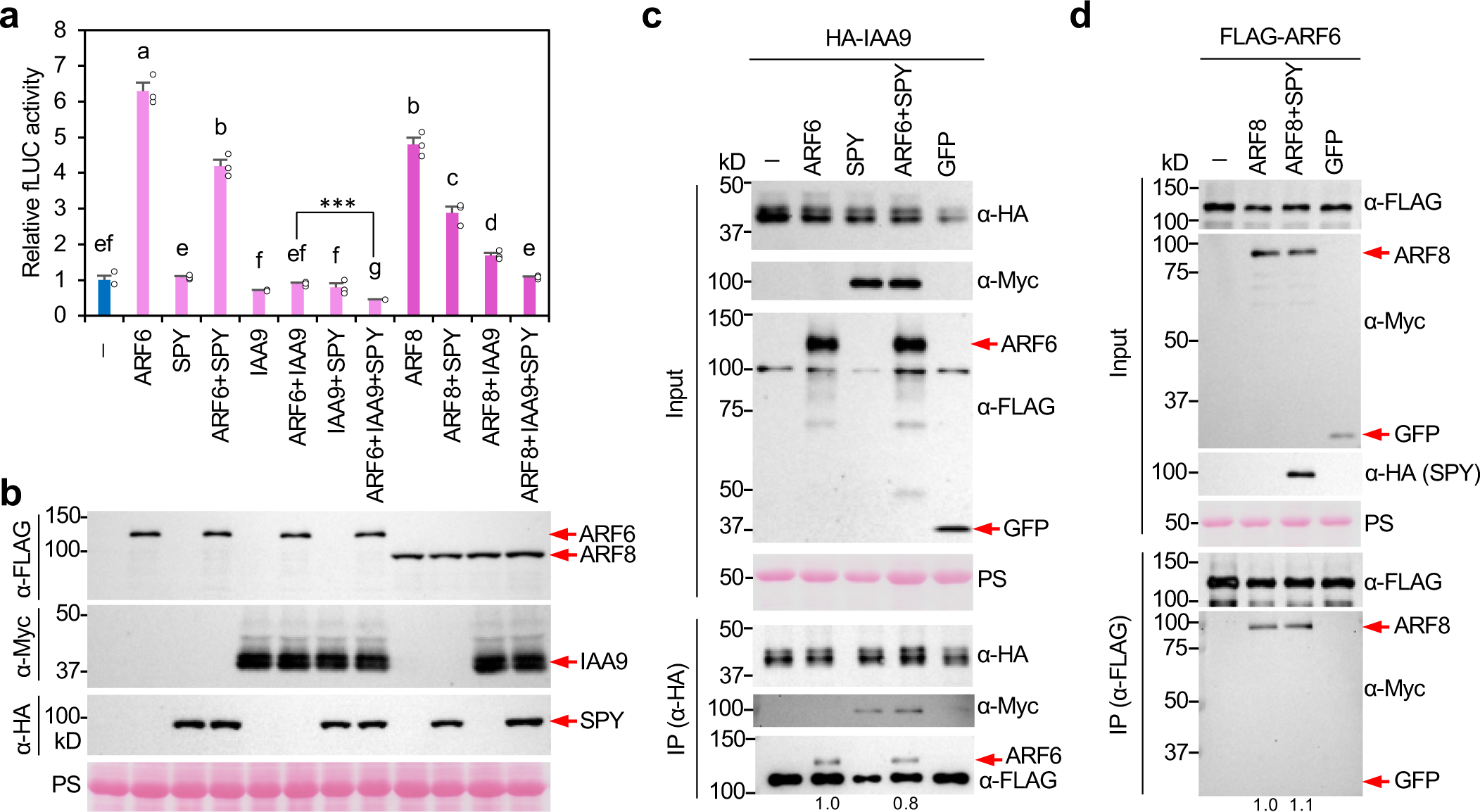
ARF6/8-IAA9 transcription repression activities were enhanced by SPY, while ARF6/8 transactivation activities were reduced by SPY. **a-b,** Dual luciferase assay in the *N. benthamiana* transient expression system showing the opposing effect of SPY on ARF vs ARF+IAA9. *35S:Renilla LUC (rLUC)* was the internal control for transformation efficiency. The reporter construct contained *P3(2x):fLUC*. Effector constructs included *35S:FLAG-ARF6*, *35S:FLAG-ARF8, 35S:Myc-IAA9^P^*^188^*^S^*, or *35S:HA-SPY* as labeled. In **a**, relative fLUC activity was calculated by normalizing with rLUC activity in each sample. Means ± SE of 3 biological replicas are shown. Different letters above the bars represent significant differences (*p* < 0.05) as determined by Tukey’s HSD mean separation test. *** *p* = 0.0002. In **b,** each effector protein was expressed at similar levels in different samples. Effector proteins in *N. benthamiana* extracts were detected by immunoblot using anti-FLAG, anti-Myc and anti-HA antibodies as labeled. **c,** Co-IP assay showing that SPY did not affect ARF6-IAA9 interaction in *N. benthamiana*. HA-IAA9^P188S^ was expressed alone or co-expressed with FLAG-ARF6, Myc-SPY or FLAG-ARF6+Myc-SPY or Myc-GFP (a negative control). Anti-HA beads were used for IP, and input and IP’ed samples were detected with anti-HA, anti-Myc and anti-FLAG antibodies, separately. FLAG-ARF6 in the IP eluate from HA-IAA9^P188S^+FLAG-ARF6 sample was set as 1.0. **d,** Co-IP assay showing that SPY did not affect ARF6-ARF8 dimerization in *N. benthamiana*. FLAG-ARF6 was expressed alone or co-expressed with Myc-ARF8 +/-HA-SPY. Myc-GFP was included as a negative control. Anti-FLAG beads were used for IP, and input and IP’ed samples were analyzed by immunoblotting with different antibodies as labeled. Myc-ARF8 in the IP eluate from FLAG-ARF6+Myc-ARF8 sample was set as 1.0. In **b-d**, PS-stained blot images showing even loading. In **a-d**, two biological repeats showed similar results.

### SPY reduced ARF6 interaction with Mediator 8

Considering that SPY enhanced the transcription repression activity of the IAA9-ARF6/8 complexes, we tested whether SPY promotes ARF6-IAA9 interaction by co-IP assay. HA-IAA9 was expressed alone or co-expressed with FLAG-ARF6 and/or Myc-SPY or FLAG-GFP (as a negative control) in *N. benthamiana*. After immunoprecipitated with anti-HA beads, we found that IAA9 binding to ARF6 was not affected by co-expression of SPY (**Fig. 6c**). Notably, IAA9 also interacted with SPY (**Fig. 6c**), and its N-terminal region (amino acid residues 1-208) was *O*-fucosylated by SPY (**Supplementary Fig. 10**). Because ARFs function as homo- and hetero-dimers^77^, we examined whether SPY affects ARF6-ARF8 dimerization by co-IP assays. FLAG-ARF6 was expressed alone or co-expressed with Myc-ARF8 and/or Myc-SPY or Myc-GFP in *N. benthamiana*. After immunoprecipitated with anti-FLAG beads, we found that SPY did not alter ARF6-ARF8 binding affinity (**Fig. 6d**). Recent studies indicate that the Mediator complex is required for class A ARF7/19-mediated transcription activation of auxin-induced genes for lateral root initiation. ARF7 and ARF19 interact directly with MED8 of the head Mediator module and MED25 of the tail Mediator module^16^. By yeast two-hybrid (Y2H) assay, we showed that both ARF6 and ARF8 interacted with MED8 (**Fig. 7a**), but not with MED25 (**Supplementary Fig. 11a**). Notably, MED8 binds to the MR fragments of ARF6 and ARF8 (**Fig. 7b**), which contain *O*-Fuc sites modified by SPY (**Fig. 3**). Moreover, AAL pulldown assay showed that MED8 was also *O*-fucosylated by SPY when transiently co-expressed in *N. benthamiana* (**Supplementary Fig. 11b**), which is consistent with the detection of several *O*-Fuc-MED8 peptides from *Arabidopsis* extracts in a recent proteomic study^35^. To test whether SPY modulates ARF-MED8 interaction, a co-IP assay was performed using *N. benthamiana* that expressed FLAG-ARF6 alone or co-expressed with Myc-MED8 and/or HA-SPY, HA-spy-19 or Myc-GFP. After IP’ed with anti-Myc beads, we found that co-expression of SPY, but not spy-19, reduced MED8 binding to ARF6 (**Fig. 7c**). To confirm this observation in *Arabidopsis*, we generated *35S:MED8-GFP* transgenic line and showed that MED8-GFP was functional in planta to rescue the *med8-2* mutant phenotype (**Supplementary Fig. 11c, 11d**). We also generated transgenic *Arabidopsis* carrying both *P_UBQ10_:FLAG-ARF6* and *35S:MED8-GFP* in WT or *spy* background. Co-IP assays were performed using protein extracts from transgenic lines carrying either *P_UBQ10_:FLAG-ARF6* alone or both *P_UBQ10_:FLAG-ARF6* and *35S:MED8-GFP* in WT or *spy* background (**Fig. 7d**). Importantly, the *spy* mutation enhanced MED8-ARF6 interaction without altering MED8 protein levels, supporting that *O*-fucosylation of ARF6 and MED8 reduced their binding affinity in planta.

**Figure 7.**
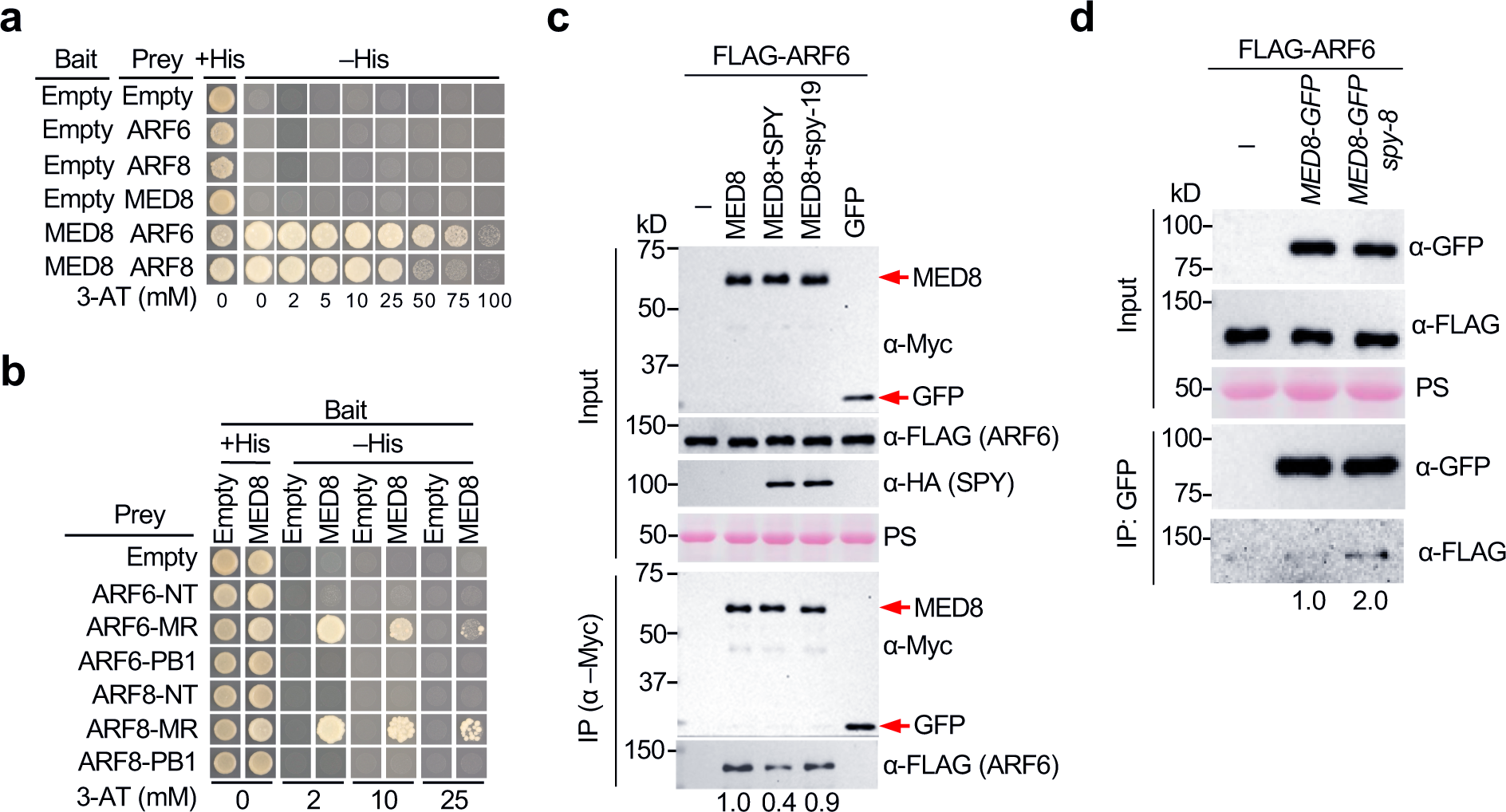
SPY reduced ARF6-MED8 interaction. **a-b,** ARF6 and ARF8 interacted with MED8 and the MR of ARFs contain the MED8 binding sequence in Y2H assay. The strength of interaction was indicated by the ability of cells to grow on –His plates with 0-100 mM 3-AT as labeled. **c,** Co-IP assay showing SPY but not spy-19 reduced ARF6-MED8 interaction in *N. benthamiana*. FLAG-ARF6 was expressed alone or co-expressed with Myc-MED8 –/+ HA-SPY, HA-spy-19 or Myc-GFP (a negative control). Anti-Myc beads were used for IP. Input and IP’ed samples were detected by immunoblot analysis as labeled. The FLAG-ARF6 protein levels in the IP eluate from FLAG-ARF6+Myc-MED8 sample was set as 1.0. **d,** Co-IP assay in *Arabidopsis* showing *spy* mutation enhanced ARF6-MED8 interaction. Transgenic lines carrying either *P_UBQ10_:FLAG-ARF6* or both *P_UBQ10_:FLAG-ARF6* and *35S:MED8-GFP* in WT or *spy-8* background were used for IP with anti-GFP beads. The input and IP’ed samples were detected by immunoblot analysis as labeled. The FLAG-ARF6 protein levels in the IP eluate from FLAG-ARF6+GFP-MED8 sample was set as 1.0. In **c-d**, PS-stained blot images showing even loading. In **a-d**, two biological repeats showed similar results.

## DISCUSSION

In this study, we demonstrated that SPY inhibits auxin-induced fruit growth in *Arabidopsis* by *O*-fucosylating ARF6/8, IAA9 and MED8. Co-IP assays showed that SPY-mediated *O*-fucosylation of these proteins reduced ARF-MED8 interaction, which leads to enhanced transcription repression activity of the ARF6/8-IAA9 complex (**Fig. 8**), presumably before pollination when auxin levels are low in the pistil. After pollination, elevated auxin levels in the pistil trigger IAA9 degradation and release ARF6 and ARF8 to activate fruit growth-related genes by interacting with MED8 of the coactivator Mediator complex (**Fig. 8**). This recruitment of the Mediator complex is known to promote the assembly of the RNA polymerase II preinitiation complex (PIC)^78,79^. SPY-mediated *O*-fucosylation of ARF6/8 and MED8 also attenuates transactivation activities of ARF6/8 by reducing ARF-MED8 interaction. We found that SPY protein levels and its POFUT activity were reduced after pollination, although the mechanism is unknown. This further enhances ARF6/8 transactivation activities by promoting ARF-MED8 interaction. Importantly, we mapped the MED8 interaction domain in ARF6/8 to their MRs, which also contain SPY target sites for *O*-fucosylation. This adds a novel regulatory mechanism via PTM to modulate ARF activity, in addition to altered binding affinity to IAA proteins by phosphorylation of DBD and/or MR of ARF5 and ARF7^29,80^, reduced DNA binding by SUMOylation in the DBD of ARF7/19^30^, and decreased protein stability by ubiquitination^31,68^. Besides *O*-fucosylation, ARF6 and ARF8 are highly phosphorylated and *O*-GlcNAcylated. However, the pistils of the *sec* mutants lacked any phenotype comparing to WT, suggesting that *O*-GlcNAcylation does not play a significant role in regulating ARF6/8 during fruit set. This could be due to a lower expression of SEC in the pistil than other tissues (Arabidopsis eFP Browser^43^).

**Figure 8.**
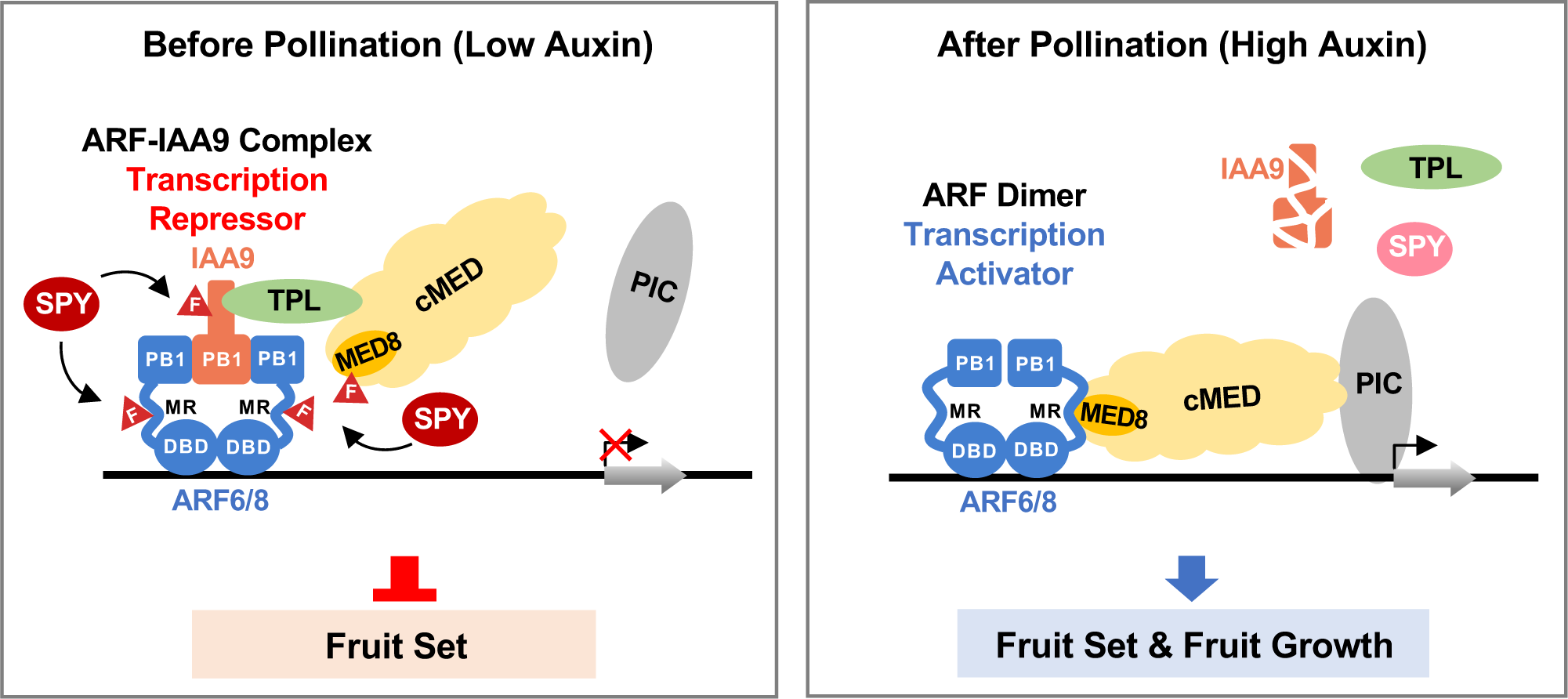
Model of regulatory mechanism of ARF activity by SPY-mediated *O*-fucosylation in fruit growth. Before pollination, the IAA9-ARF6/8 complexes function as transcription repressors to inhibit fruit set. SPY *O*-fucosylates ARF6/8, IAA9 and MED8, a subunit of the core Mediator complex (cMED), which reduces ARF-MED8 interaction to enhance transcription repression activities of the ARF-IAA9 complexes. The co-repressor TPL, recruited by IAA9, also interferes with ARF binding to cMED^15^ and may recruit the CKM repressive module (not shown) to block transcription of ARF target genes^16^. After pollination, elevated auxin levels in the pistil trigger IAA9 degradation and release ARF6 and ARF8 homo and/or hetero-dimers to activate fruit growth-related genes by recruiting the coactivator Mediator complex and promoting the assembly of RNA Pol II preinitiation complex (PIC). In addition, SPY protein level and/or activity are reduced after pollination via an unknown mechanism. This further enhances ARF6/8 transactivation activities by promoting ARF-MED8 interaction.

Although recent *O*-fucosylome studies have identified hundreds of SPY target proteins^34,35^, the molecular mechanism of SPY regulation has only been elucidated for a handful of its targets. These include enhanced protein-protein interaction for GA signaling repressors DELLAs^32^, increased or decreased protein stability for bHLH transcription factors TEOSINTE BRANCHED 1, CYCLOIDEA, PCFs (TCP14/15) in cytokinin response, and transcription repressor PSEUDO RESPONSE REGULATOR 5 in circadian clock, respectively^34,37,39^, DNA binding and/or transcription repression activity of bHLH transcription factor SPATULA for style development ^38^, and protein translocation of co-chaperonin CPN20 in abscisic acid signaling^36^. Here, we found that SPY not only *O*-fucosylates ARF6 and ARF8, but also two ARF6/8-interacting proteins, IAA9 and MED8, resulting in reduced interaction between ARF and MED8. Thus, SPY-mediated *O*-fucosylation of multiple components in nuclear protein complexes may contribute to another layer of transcription regulation.

The MRs of class A ARFs are enriched in glutamine residues and contain intrinsically disordered regions (IDRs)^18,19,67^. The MR, together with the PB1 domain, of ARF7 and ARF19, are required for cytoplasmic condensate formation, which blocks their function by preventing nuclear localization^67^. Intriguingly, unlike ARF7 and ARF19, we did not observe cytoplasmic condensate formation of ARF6 in the root cells of the maturation zone. The ARF6-GFP fusion protein was only detected in the nucleus, suggesting that nucleocytoplasmic partitioning may not be a universal regulatory mechanism for all class A ARFs. Previous studies also showed that the MR of different class A ARFs specifies the distinct function of individual ARF in transcription regulation by interactions with chromatin remodelers^81^ and/or transcription factors^18^. Here we show that MR of ARF6/ARF8 interacts with MED8 of the coactivator Mediator complex, and that *O*-fucosylation by SPY plays an important role in modulating this interaction. Further studies will determine whether this is a general regulatory mechanism for all class A ARFs. Besides MED8, three other MEDs (MED12, 13,14) in the Mediator complex are present in the recently reported *O*-fucosylome in *Arabidopsis*^35^. Our study lays the foundation for elucidating the molecular mechanism of SPY-mediated transcription regulation through its modulation of the coactivator MED complex. Finally, understanding the regulatory mechanism of class A-ARFs-controlled parthenocarpy has significant implication in developing climate-resilient crops as production of seedless fruit without fertilization can ensure consistent fruit yield under stressful environmental conditions.

## METHODS

### Plant materials, growth conditions, plant transformation, and statistical analysis

*Arabidopsis thaliana* Columbia-0 (Col-0) was the wild-type control for most of the mutants used in this study. The exceptions are *spy-8* and *spy-19*, which are in the Landsberg *erecta* (L*er*) background. Plants were grown under long-day (22 °C, 16 h light/8 h dark) conditions. All genotyping primers used in this study are listed in **Supplementary Table 7**. The genotyping primers for *spy-3* were designed using dCAPS Finder 2.0 (http://helix.wustl.edu/dcaps/)^82^. The following mutants and transgenic lines have been described previously: (1) the *spy* mutants, *spy-3*, *spy-8*, *spy-19*, and *spy-23* (WiscDsLox241C03)^32,35,83^, and the *sec* mutants, *sec-2*^84^ and *sec-5*^85^; (2) The *arf* mutants, *arf6-2*, *nph4-1* (*arf7*), *arf8-*3, *arf6-2 nph4-*1, *nph4-1 arf8-3*, *arf6-2 -/- arf8-3 +/-* ^86–89^; (3) The *bZIP1* mutants, *bzip1-1* (SALK_059343) and *bzip1-2* (SALK_069489)^90^; (4) *P_SPY_:GFP-SPY spy-3*, *P_SPY_:GFP-SPY-NES spy-*3, and *P_SPY_:GFP-SPY-NLS spy-3*^37^; (5) *P_ARF6_:ARF6-GFP* and *P_ARF7_:ARF7-YFP* lines in the Col-0 background^67,91^; (6) *iaa9-1*^71^ and *med8-2* (CS16505)^92^ that were obtained from the Arabidopsis Biological Resource Center. For the genetic interaction study, *arf6-2, arf8-3, arf6-2 -/- arf8-3 +/-* were crossed to *spy-3*, and the F2 plants were genotyped by PCR to identify homozygous *arf spy* double mutants and *arf* sesquimants (*arf6 -/- arf8 +/-* and *arf8 -/- arf6 +/-*) in the *spy-3* background. Because the homozygous *arf6 arf8* double mutant is sterile^87^, the *arf* sesquimutants (*arf6 -/- arf8 +/-* and *arf8 -/- arf6 +/-*) in *SPY* or *spy-3* background were identified in the segregating F3 population by genotyping for phenotype analysis.

P_ARF6:_FLAG-ARF6 and P_ARF8_:FLAG-ARF8 constructs were introduced into *arf6-2* or *arf8-3* separately by Agrobacterium-mediated transformation. Homozygous transgenic lines containing a single insertion site were obtained as described previously^93^. A representative *P_ARF6:_FLAG-ARF6* line (#1-11-7) and a *P_ARF8_:FLAG-ARF8* line (#2-1-4) were used for the complementation test. P_UBQ10_:FLAG-ARF6 and P_UBQ10_:FLAG-ARF8 constructs were introduced into WT (L*er*) separately by Agrobacterium-mediated transformation. The transgenes, *P_UBQ10_:FLAG-ARF6* (in line #4-2-1) and *P_UBQ10_:FLAG-ARF8* (in line #4-1-1), were then introduced into the *spy-8* background by genetic crosses. The resulting homozygous *P_UBQ10_:FLAG-ARF6 spy-8* line and *P_UBQ10_:FLAG-ARF8 spy-8* line were obtained by genotyping. To test the activity of 35S:MED8-GFP in planta, the pGWB405:MED8-GFP construct was introduced into *med8-2* by Agrobacterium-mediated transformation. Homozygous transgenic lines containing a single insertion site were obtained, and a representative *35S:MED8-GFP med8-2* line (#1-32-1) was used for phenotype analysis. To generate the double transgenic lines for co-IP assays, *P_UBQ10_:FLAG-ARF6 spy-8* was transformed with the 35S:MED8-GFP construct (pGWB405:MED8-GFP). A representative double transgenic line *FLAG-ARF6 MED8-GFP spy-8* (#1-3-1) was crossed with *P_UBQ10_:FLAG-ARF6* in the WT (L*er*) background to generate *FLAG-ARF6 MED8-GFP* in WT (#1-4-52). The *P_ARF6_:ARF6-GFP* line in the Col-0 background was crossed with the *spy-3* mutant to generate *P_ARF6_:ARF6-GFP spy-3* line. The *P_SPY_:FLAG-SPY* construct was introduced into *spy-3* by Agrobacterium-mediated transformation to obtain the *P_SPY_:FLAG-SPY spy-3* lines, and a representative homozygous line (#1-2-17) was used for further analysis.

For agroinfiltration, 3 or 4-week-old plants of *Nicotiana benthamiana* were used. Statistical analyses were performed by Tukey’s honestly significant difference (HSD) mean separation tests with SPSS Statistics 17.0 software.

### Plasmid construction

The following plasmids were described previously: 35S:Myc-SPY and 35S:Myc-SEC for transient expression^46^; P_35S_:FLAG-GFP-NLS for the negative control of transient expression^34^; pEarleyGate201 and 203 vectors^94^, pDONR207:SPY^95^, pEG3F-GW^34^, pDEST32-HA^96^ and pDEST22-FLAG^96^ for cloning new constructs. Primers and plasmid constructs are listed in **Supplementary Tables 7 and 8**, respectively. All DNA constructs generated from PCR amplification were sequenced to ensure that no mutations were introduced.

### Flower emasculation and hormone treatments

Flower emasculation and hormone treatments were conducted following the methods described previously^42^. Flowers were emasculated at stage 10^97^, approximately 2 days before anthesis (–2 DAA), to remove sepals, petals, and anthers. The pistils of emasculated flowers were immersed uniformly in 100 µM GA_3_ or 50 µM picloram (Sigma-Aldrich, CAS-1918-02-1) solution that also contained 0.01% (v/v) Triton X-100, or in 0.01% (v/v) Triton X-100 alone for mock treatment. Seven days after emasculation, the final length of the pistils was measured using ImageJ.

### Transient expression and dual luciferase assay in *Nicotiana benthamiana*

For dual luciferase assays and pulldown assays, transient expression of different epitope-tagged proteins in *N. benthamiana* was performed as described with slight modifications^98^. The *N. benthamiana* leaves were harvest after 48 hr of agroinfiltration^99^ and luciferase activity was quantified using the dual-luciferase reporter assay system (Promega). To determine the relative promoter activity, the ratio of fLUC to rLUC activity was calculated for each sample. Three biological repeats were conducted for each effector combination.

### AAL-agarose pull-down, co-IP assays and protein blot analyses

For AAL pulldown assays, FLAG-GFP/ARF6/ARF6-N/ARF6-MR/ARF6-C or IAA9/IAA9-N/IAA9-C were individually expressed or co-expressed with Myc-SPY in *N. benthamiana* by agroinfiltration, following the established protocol^95^. Protein extracts were incubated with AAL-agarose beads (Vector Labs, AL-1393-2, 2 mg lectin/mL) to enrich for *O*-fucosylated proteins, and analyzed by immunoblot analysis as described^34^.

To investigate the effect of SPY on ARF6-IAA9 interaction, HA-IAA9^P188S^ was expressed alone or co-expressed with FLAG-ARF6 and/or Myc-SPY or FLAG-GFP in *N. benthamiana*. Co-immunoprecipitation (co-IP) assays were performed using anti-HA beads (Sigma-Aldrich, A2095), following the described procedure^15^. To examine the effect of SPY on ARF6-MED8 interaction, FLAG-ARF6 was expressed alone or co-expressed with Myc-MED8 and/or HA-SPY, or Myc-GFP *in N. benthamiana*. Co-immunoprecipitation (co-IP) assays were performed using anti-Myc beads (Sigma-Aldrich, A7470), following the described procedure^15^. To verify the effect of *spy* on ARF6-MED8 interaction in planta, co-IP assays were performed using ChromoTek GFP-Trap® Magnetic Agarose (Proteintech, GTMA-400) and protein extracts from transgenic *Arabidopsis* lines carrying either *FLAG-ARF6* or both *FLAG-ARF6* and *MED8-GFP* in WT (L*er*) or *spy-8* background.

Immunoblot analyses were performed using horseradish peroxidase (HRP)-conjugated anti-FLAG M2 mouse monoclonal (Sigma-Aldrich A8592, 1:10,000 dilution) and mouse HRP-anti-MYC monoclonal antibodies (BioLegend #626803, 1:5,000 dilution), mouse HRP-anti-HA (6E2) monoclonal antibody (Cell Signaling Technology #2999S, 1:5,000 dilution), mouse anti-GFP (Roche #11814460001, 1:2,000 dilution), mouse anti-Tubulin (Sigma T5168, dilution 1:500,000). HRP-conjugated donkey anti-mouse IgG (Jackson ImmunoResearch #715-035-150) was used to detect anti-GFP at 1:5,000 dilution. Biotinylated-Aleuria aurantia lectin (AAL-biotin, Vector Labs B-1395, 1:30,000) followed by Streptavidin-HRP (Jackson ImmunoResearch #016-030-084, 1: 100,000x dilution,) were used for AAL blot. To detect *O*-GlcNAcylated proteins, the anti-*O*-GlcNAc monoclonal antibody CTD110.6 (5,000× dilution, Cell Signaling Technology, Cat. No. 9875) followed by a goat anti-mouse IgM-HRP secondary antibody (30,000× dilution, Thermo-Fisher Scientific, Cat. No. 31440) were used.

### Y2H assays

The ProQuest Two-Hybrid system (Invitrogen) and yeast strain pJ69-4A were used for the Y2H assays. Yeast transformation and 3-amino-1,2,4-triazole (3-AT) tests were performed according to the manufacturer’s protocol as described previously^96^ with slight modifications: 3-AT concentrations in the plates were 0, 2, 5, 10, 25, 50, 75, and 100 mM in most cases, except that 0, 2, 10, 25 mM were used for the ARF6 or ARF8 domain mapping.

### Confocal microscopy

GFP signals in pistils were analyzed using *P_SPY_:GFP-SPY spy-3*, *P_SPY_:GFP-SPY-NES spy-*3, and *P_SPY_:GFP-SPY-NLS spy-3*^37^. The pistils at stage 10 were imaged using a Zeiss 880 equipped with a 20x objective. The pistils from stage 10 to stage 14 were imaged using a Zeiss Axio Zoom.V16. Identical image settings were used for direct comparison.

For detecting the protein localization of ARF6-GFP and ARF7-YFP in the WT or *spy-3* background, the primary root cells in 3d-old seedlings were stained with 10 μM propidium iodide (Sigma, P4170) and imaged using a Zeiss 880 equipped with a 20x objective. The pistils at stage 10 were also analyzed. Excitation and detection were set as follows: GFP or YFP, excitation at 488 nm and detection at 493–558 nm; PI staining, excitation at 561 nm and detection at 605–695 nm. Confocal images were processed using the Fiji package of ImageJ. Identical image settings were used for direct comparison.

### RNA-seq analysis

Total RNAs were purified from unpollinated pistils from stage 10 flowers (–2 DAA) of *arf6-2, arf8-3, spy-3* and WT (Col-0), and pollinated pistils from WT stage 14 flowers (0 DAA). RNA-Seq cDNA libraries (two biological repeats) were prepared with the QuantSeq 3’mRNA-Seq library prep kit FWD for Illumina (Lexogen). DNA sequencing was performed with Illumina Next-Seq500 High-Output 75bp SR. Sequence alignment and DE (differential expression) analysis were conducted on a commercial server with pre-established computational pipelines (8 omics Gene Technology Co. Ltd., Beijing, China)^100^. Co-regulated genes among *spy-3*, *arf6-2*, *arf8-3* and fertilization-responsive gene lists were then identified (fold change > 1.5; *p* < 0.05). Venn diagrams were made using online tool at InteractiVenn.net^101^. GO analysis was performed using Panther v.18.0^102^. Heatmap analysis was made by using online tool at MetaboAnalyst 5.0^103^.

### RT-qPCR and ChIP-qPCR analyses

Total RNAs from pistils stage 10 or stage14 were isolated with RNeasy Plant Mini Kit (Qiagen). First-strand cDNA was then synthesized using a Transcriptor First Strand cDNA Synthesis kit (Roche). qPCR analyses were performed using FastStart Essential DNA Green Master mix (Roche) and LightCycler 96 instrument (Roche). The PCR program was performed as described before^96^. Relative transcript levels were determined by normalizing with *PP2A* (At1g13320). Mean values of fold change were calculated from three biological replicates. For ChIP-qPCR analysis, seedlings of the *P_UBQ10_:FLAG-ARF6* and *P_UBQ10_:FLAG-ARF8* transgenic lines in the WT or *spy-8* background were grown in liquid 0.5x MS and 1% sucrose for 10 days in continuous light. The ChIP procedure was performed as described^104^. Primers for all qPCR analyses are listed in **Supplementary Table 7**.

### Tandem affinity purification of FLAG-ARF6/8 from *N. benthamiana* or *Arabidopsis* for immunoblot or MS analysis

The FLAG-ARF6/8 proteins (containing a 6xHis-3xFLAG-tag) that were transiently expressed in *N. benthamiana* were tandem affinity-purified using a His-Bind resin followed by anti-Flag-M2-agarose beads (Sigma-Aldrich) as described^32^ with slight modifications. 3 g of starting tissue was used for MS analysis and a smaller scale with 0.7 g of starting tissue was used for AAL blot analysis. The extraction and purification buffer included 50 mM fucose and 2x protease inhibitors. His-FLAG-ARF6/8 proteins were also purified from *P_UBQ10_:FLAG-ARF6/8* transgenic *Arabidopsis* lines in WT or *spy-8* backgrounds. The tandem affinity purification procedures were as previously described^32^ for both AAL blot and MS analyses, with the following modifications: 10 g of starting tissues was used, and 50 mM fucose and 2x protease inhibitors were included during purification. The cleared extract was incubated with 0.4 mL of His-Bind resin for 1.5 h at 4 °C and was loaded onto a disposable plastic column. The second purification step was carried out with 10 μL of anti-Flag-M2-agarose beads (Sigma-Aldrich). The purified protein was digested by trypsin on-beads for MS analysis as described^32^.

### Identification of *O-*fucosylation sites by liquid chromatography (LC)–electrospray ionization (ESI)–mass spectrometry (MS)

Trypsin-digested 6His-3xFLAG-ARF6 and 6His-3xFLAG-ARF8 proteins, purified from *N. benthamiana* and from *A. thaliana*, were separated by reverse phase nano-HPLC, and analyzed by electrospray ionization (ESI)-MS using a Thermo Scientific^TM^ Orbitrap Fusion^TM^ Tribrid^TM^ mass spectrometer equipped with electron transfer dissociation (ETD)^105^. Nano-HPLC was performed as described previously^32^. MS1 spectra were acquired in the Orbitrap at a resolution of 60,000 followed by data-dependent, 3 second Top-N, MS/MS experiments. Precursors were isolated by resolving quadrupole with a 3 m/z window. An MS/MS decision tree was made for each sample that included collision-activated dissociation CAD, ETD, and higher-energy collisional dissociation (HCD). Precursors with a charge state less than 5 were selected for low resolution MS/MS, fragmented with CAD and ETD, and scanned out of the linear ion trap at a normal scan rate. Calibrated reaction times were used for ETD events. Precursors with a charge state above 4 were selected for high resolution MS/MS, fragmented with HCD and ETD, and scanned out of the Orbitrap at a resolution of 30,000. Electron transfer-higher energy collisional dissociation (EThcD) was later added to the decision tree for precursors that had a charge state above 4 and m/z above 900, and scanned out of the Orbitrap at a resolution of 30,000. For some samples, peptides were targeted for fragmentation based on intact mass and retention time observed in previous experiments.

To detect additional ARF6 peptides within the MR sequence, trypsin-digested ARF6 samples from *N. benthamiana* underwent a second digestion with chymotrypsin overnight at room temperature. ARF6 and ARF8 samples from *N. benthamiana* were subsequently cleaned up using hydrophilic interaction liquid chromatography (HILIC) prior to a second MS analysis, a method developed by Keira Mahoney (unpublished data) and previously described^34^. Cleaned up samples were analyzed immediately by MS or stored at -35°C.

Data files were searched using Byonic Version 3.8.13 (Protein Metrics)^106^. ARF6 and ARF8 data files were searched against a database containing the sequence of 6His-3xFLAG-ARF6 and 6His-3xFLAG-ARF8, respectively. Search parameters included specific cleavage C-terminal to R and K residues with up to five allowed missed cleavages, 10 ppm tolerance for precursor mass, 15 ppm tolerance for high-resolution MS/MS, and 0.35 Da mass tolerance for low-resolution MS/MS. After the chymotrypsin digest, cleavages C-terminal to F, L, Y, and W residues were added to the search parameters. Variable modifications selected included oxidation of M residues, phosphorylation of S, T, and Y residues, alkylation of C residues, and *O*-GlcNAcylation, *O*-fucosylation, and *O*-hexosylation of S and T residues. No protein false discovery rate cutoff or score cutoff was applied prior to the output of search results. Peptides were manually validated, and modification sites were determined manually using ETD spectra.

### Accession Numbers

Arabidopsis Genome Initiative locus identifiers for the genes mentioned in this article are as follows: *SPY* (AT3G11540), *ARF6* (AT1G30330), *ARF7* (*NPH4*, AT5G20730), *ARF8* (AT5G37020), *IAA9* (AT5G65670), *SEC* (AT3G04240), *MED8* (AT2G03070), *MED25* (AT1G25540), *PP2A* (AT1G13320), *ERF023* (AT1G01250), *ERF107* (AT5G61590), *bZIP1* (AT5G49450), *AGP12* (AT3G13520), *SHL* (AT4G39100), *GNTL* (AT3G52060), *IAA19* (AT3G15540).

## Supporting information

Supplementary Data Sets 1-4

## Data Availability

Raw and processed RNA-Seq data have been deposited at in the NCBI Sequence Read Archive under BioProject PRJNA1095421. The mass spectrometry proteomics data have been deposited to the ProteomeXchange Consortium via the PRIDE^107^ partner repository with the dataset identifier PXD051232 (Project DOI: 10.6019/PXD051232). Source Data files will be provided with this paper before publication.

## Acknowledgements

We thank Jason Reed and Lucia Strader for helpful discussions, and for sharing *Arabidopsis arf* mutants and reporter lines. We also thank Jyan-Chyun Jang for providing the *bzip1* mutants. We are grateful to Mingyuan Zhu for helping with confocal microscopy analysis. This work was supported by the National Institutes of Health (GM100051 and GM150029 to TPS, GM037537 to DFH), and the United State Department of Agriculture (2023-67013-39532 to TPS). A special thank you to Protein Metrics for providing Byonic.

## Author Contributions

T.P.S. and Y.W. conceived and designed the research project. Y.W., R.Z. and J.H. performed experiments, and T.P.S., Y.W., R.Z. and J.H. analyzed the data and generated figures. L.W. and H.W. helped with RNA-seq data analysis. S.K. performed LC-ESI-MS analysis, and S.K., J.S., and D.F.H. analyzed MS data. T.P.S. wrote the manuscript with input from all co-authors.

## FIGURE LEGENDS

**Supplementary Figure 1. SPY expression patterns and levels in pistils. a,** Confocal microscopy showing localization of GFP-SPY in both cytoplasm and nucleus, GFP-SPY-NES in the cytoplasm, and GFP-SPY-NLS in the nucleus. Images of pistils at 2 days before anthesis (–2 DAA). Bar = 20 µm. **b,** GFP-SPY, GFP-SPY-NLS and GFP-SPY-NES were accumulated at similar levels in these transgenic lines. Protein blot was probed with an anti-GFP antibody. **c,** FLAG-SPY was reduced in the *P_SPY_:FLAG-SPY spy-3* line after anthesis. **d,** *P_SPY_:FLAG-SPY spy-3* line showed reduced protein *O*-fucosylation at 3 DAA and 5 DAA compared to that at –2 DAA and 0 DAA. O-fucosylated proteins in total proteins extracted from the *P_SPY_:FLAG-SPY spy-3* line and *spy-3* (a negative control) were detected by protein blot analysis using AAL-biotin. * indicates reduced *O*-fucosylated proteins. In **b-d**, Ponceau S (PS)-stained blot showing protein loading. In **a-d**, two biological repeats showed similar results.

**Supplementary Figure 2. SPY-regulated fruit growth is mediated by ARF8 together with ARF6, but not ARF7 (NPH4). a,** Pistil width of *spy-3*, *arf6-2* and *arf8-3* mutants. Mutant alleles are homozygous unless specified as heterozygous, including arf6+/- and arf8+/-. **b-c**, Epistasis analysis of *arf6, nph4-1* and *arf8*. The *nph4-1* mutation did not display parthenocarpy after emasculation. In **b**, photo showing representative pistils of different genotypes 7 days after emasculation. Bar = 2 mm. In **c**, pistil lengths, n>15. In boxplots **a** and **c**, center lines and box edges are medians and the lower/upper quartiles, respectively. Whiskers extend to the lowest and highest data points within 1.5x interquartile range (IQR) below and above the lower and upper quartiles, respectively. Different letters above the boxes represent significant differences (*p* < 0.05) as determined by Tukey’s HSD mean separation test. In **a-c**, two biological repeats showed similar results.

**Supplementary Figure 3. FLAG-ARF6/8 partially rescued the *arf6* and *arf8* parthenocarpic phenotype. a,** photo showing representative pistils of different lines 7 days after emasculation. Bar = 2 mm. **b**, pistil lengths. n>15. The center lines and box edges in the box plot are medians and the lower/upper quartiles, respectively. Whiskers extend to the lowest and highest data points within 1.5x IQR below and above the lower and upper quartiles, respectively. Different letters above the boxes represent significant differences (*p* < 0.05) as determined by Tukey’s HSD mean separation test. Two biological repeats showed similar results.

**Supplementary Figure 4. ARF6 and ARF8 are *O*-fucosylated and *O*-GlcNAcylated. a-b,** FLAG-ARF6 and -ARF8 were *O*-fucosylated by SPY and *O*-GlcNAcylated by SEC, respectively. FLAG-ARF6/8 proteins were expressed alone or co-expressed with SPY or SEC in *N. benthamiana*. Immunoblots containing affinity-purified FLAG-ARF6/8 were probed with anti-FLAG, AAL-biotin or anti-*O*-GlcNAc antibody as labeled. Arrow in the top panel indicates FLAG-ARF6 or -ARF8, arrow in the middle panel indicates *O*-fucosylated FLAG-ARF6 or - ARF8, and arrow in the bottom panel indicates *O*-GlcNAcylated FLAG-ARF6 or -ARF8. **c-d,** *sec* mutants did not alter growth of unpollinated pistils. **c,** photo showing representative pistils of different genotypes 7 days after emasculation. Bar = 1 mm. **d**, pistil lengths. n>15. The center lines and box edges in the box plot are medians and the lower/upper quartiles, respectively. Whiskers extend to the lowest and highest data points within 1.5x IQR below and above the lower and upper quartiles, respectively. Different letters above the boxes represent significant differences (*p* < 0.05) as determined by Tukey’s HSD mean separation test. Two biological repeats showed similar results.

**Supplementary Figure 5. PTM sites in ARF6 and ARF8. a-b,** *O*-Fuc, O-GlcNAc and phosphorylation sites in ARF6 and ARF8 identified by MS analysis. The schematic shows the ARF6 (**a**) or ARF8 protein (**b**); The marked S/T residues are confirmed PTM sites. The sequence within brackets contains PTM(s) for which the specific residue(s) could not be determined. * indicates PTM that was reported previously^35,47^.

**Supplementary Figure 6. Identification of coregulated genes among fertilization-responsive vs ARF6-, ARF8- and SPY-responsive genes in pistils by RNA-seq analysis.** RNA-seq analysis was performed using −2 DAA pistils of *arf6, arf8, spy-3* and WT, and 0 DAA WT pistils. The differentially expressed gene (DEG) lists for ARF6-, ARF8- and SPY-responsive genes, and for fertilization-responsive genes (WT 0 DAA vs WT –2 DAA) are in **Supplementary Table 2**. **a,** Total, up- or down-regulated DEGs in WT 0 DAA or in each mutant (vs WT –2 DAA). **b-d.** Venn diagrams of coregulated DEGs among fertilization (WT 0 DAA vs WT –2 DAA), *arf6, arf8* and *spy-3*. **e**, Heat map of coregulated genes among fertilization-, ARF8- and SPY-responsive DEGs. **f**, Venn diagram of overlapping genes between total SPY/ARF8 coregulated genes (181 DEGs) and the ARF6 ChIP-seq gene list^51^.

**Supplementary Figure 7. The *bzip1* and *iaa9* mutants displayed longer pistils after emasculation.** In **a** and **c**, photos showing representative pistils of different genotypes 7 days after emasculation. Bar = 2 mm. In **b** and **d**, pistil lengths were shown in boxplots. n>15. The center lines and box edges are medians and the lower/upper quartiles, respectively. Whiskers extend to the lowest and highest data points within 1.5x IQR below and above the lower and upper quartiles, respectively. Different letters (in **b**) or the asterisk (in **d**) above the boxes represent significant differences (*p* < 0.05) as determined by Tukey’s HSD mean separation test. Two biological repeats showed similar results.

**Supplementary Figure 8. ARF6 and ARF8 protein accumulation or nuclear localization were not affected by *spy*. a**, *P_UBQ10_*:*FLAG-ARF6* and *P_UBQ10_*:*FLAG-ARF8* in WT vs *spy* background. Immunoblots containing total proteins extracted from seedlings were probed with anti-FLAG or anti-tubulin antibody. **b**-**d**, *P_ARF6_*:*ARF6-GFP* in WT vs *spy* background. In **c**, *P_ARF7_*:*ARF7-YFP* in WT background was included as a control. GFP/YFP signals detected by confocal microscopy showing root tips (**b**) or upper roots (**c**) of 3d-old seedlings or stage-10 pistils (**d**). In **c,** ARF6-GFP was only detected in the nuclei of root cells in the maturation zone, whereas ARF7-YFP localized in cytoplasmic condensates. In **b-c**, roots were stained with propidium iodide before imaging. In **b-d**, bar = 10 µm.

**Supplementary Figure 9. ARF6 transactivation activities were reduced by SPY, but not by spy-19. a-b,** Dual luciferase assay in the *N. benthamiana* transient expression system. *35S:Renilla LUC (rLUC)* was the internal control for transformation efficiency. The reporter construct contained *P3(2x):fLUC*. Effector constructs included *35S:FLAG-ARF6* and/or *35S:HA-SPY,* and/or *35S:HA-spy-19* as labeled. In **a**, relative fLUC activity was calculated by normalizing with rLUC activity in each sample. Means ± SE of 3 biological replicas are shown. Different letters above the bars represent significant differences (*p* < 0.05) as determined by Tukey’s HSD mean separation test. In **b,** each effector protein was expressed at similar levels in different samples. Effector proteins in *N. benthamiana* extracts were detected by immunoblot using anti-FLAG, anti-Myc and anti-HA antibodies as labeled. Ponceau S (PS)-stained gel images showing similar sample loading. Two biological repeats showed similar results.

**Supplementary Figure 10. AAL pulldown assay showed that N-terminal domain of IAA9 was *O*-fucosylated by SPY.** FLAG-tagged full-length (FL) or truncated IAA9 proteins were expressed alone (–) or co-expressed (+) with Myc-SPY in *N. benthamiana.* FLAG-GFP, a negative control. *O*-fucosylated proteins were pull-downed by AAL-agarose. Immunoblot containing input (top panel) or AAL-agarose pull-down samples (bottom panel) was probed with anti-FLAG and anti-Myc antibodies as labeled. PS, Ponceau S-stained blot showing even loading. N, N-terminal domain of IAA9 (amino acid residues 1-208); C, C-terminal PB1 domain of IAA9 (amino acid residues 209-326). Two biological repeats showed similar results.

**Supplementary Figure 11. SPY *O*-fucosylates MED8. a,** ARF6/8 did not interact with MED25 in Y2H assay. **b,** MED8 was *O*-fucosylated by SPY. FLAG-MED8 was expressed alone (–) or co-expressed (+) with SPY in *N. benthamiana*. Immunoblots containing affinity-purified FLAG-MED8 proteins were probed with AAL-biotin or anti-FLAG antibody as labeled. **c-d,** The *35S:MED8-GFP* transgene rescued the late flowering phenotype of *med8-2*. In **c**, photo was taken at 30d-old, and bar = 2 cm. In **d**, n=10. The center lines and box edges in the box plot are medians and the lower/upper quartiles, respectively. Whiskers extend to the lowest and highest data points within 1.5x IQR below and above the lower and upper quartiles, respectively. Different letters above the boxes represent significant differences (*p* < 0.05) as determined by Tukey’s HSD mean separation test. Two biological repeats showed similar results.

**Supplementary Table 1. Summary of ARF6 and ARF8 PTMs by MS analysis**

**Supplementary Table 2. RNA-seq DEGs**

**Supplementary Table 3. Co-regulated gene lists**

**Supplementary Table 4. GO term analysis of SPY/ARF8 coregulated genes**

**Supplementary Table 5. Selected genes for RT-qPCR and ChIP-qPCR**

**Supplementary Table 6. Overlap between SPY/ARF8-DEGs and ARF6 ChIP-seq dataset**

**Supplementary Table 7. List of primers**

**Supplementary Table 8. List of constructs**

