## Supplementary Figures for "The *O*-Fucosyltransferase SPINDLY Attenuates Auxin-Induced Fruit Growth by Inhibiting ARF6 and ARF8 binding to Coactivator Mediator Complex in *Arabidopsis*"

Supplementary Figure 1

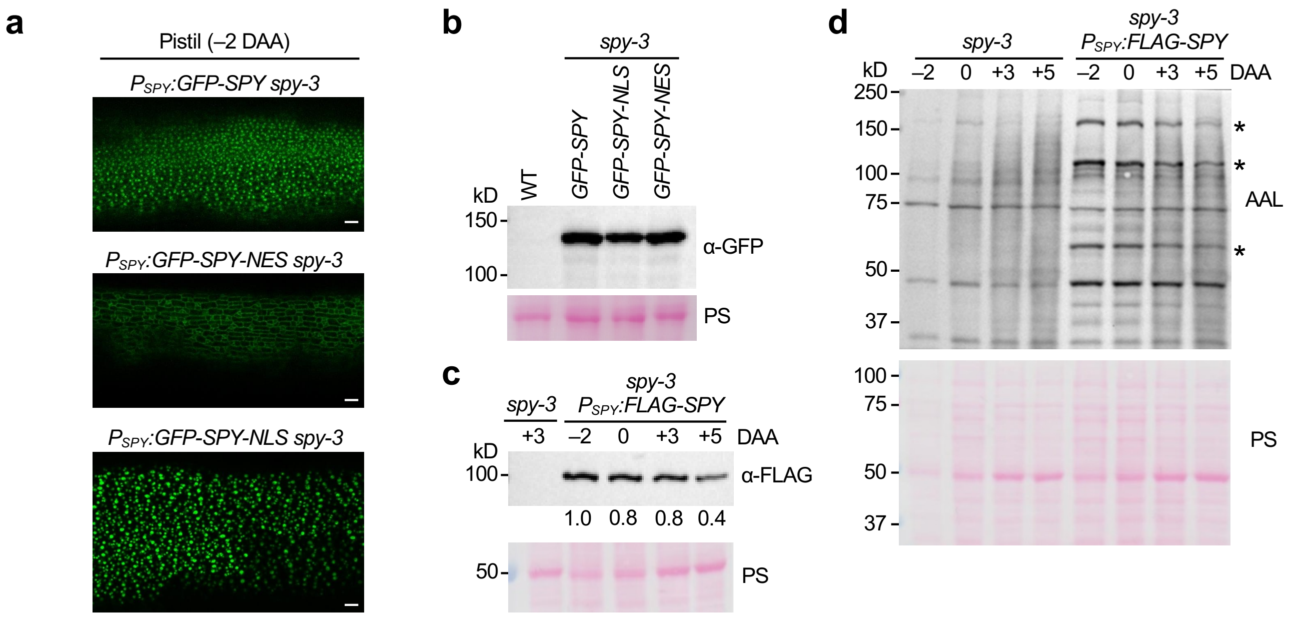


**Supplementary Figure 1. SPY expression patterns and levels in pistils. a,** Confocal microscopy showing localization of GFP-SPY in both cytoplasm and nucleus, GFP-SPY-NES in the cytoplasm, and GFP-SPY-NLS in the nucleus. Images of pistils at 2 days before anthesis (–2 DAA). Bar = 20 µm. **b,** GFP-SPY, GFP-SPY-NLS and GFP-SPY-NES were accumulated at similar levels in these transgenic lines. Protein blot was probed with an anti-GFP antibody. **c,** FLAG-SPY was reduced in the $P_{SPY}$*:FLAG-SPY spy-3* line after anthesis. **d,** $P_{SPY}$*:FLAG-SPY spy-3* line showed reduced protein *O*-fucosylation at 3 DAA and 5 DAA compared to that at –2 DAA and 0 DAA. O-fucosylated proteins in total proteins extracted from the $P_{SPY}$*:FLAG-SPY spy-3* line and *spy-3* (a negative control) were detected by protein blot analysis using AAL-biotin. * indicates reduced *O*-fucosylated proteins. In **b-d**, Ponceau S (PS)-stained blot showing protein loading. In **a-d**, two biological repeats showed similar results.

Supplementary Figure 2

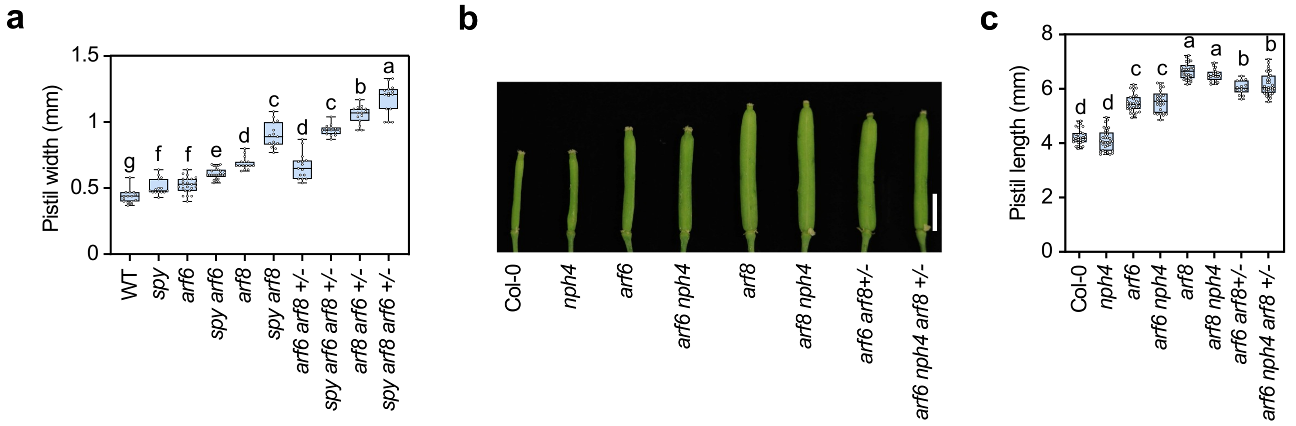


**Supplementary Figure 2. SPY-regulated fruit growth is mediated by ARF8 together with ARF6, but not ARF7 (NPH4). a,** Pistil width of *spy-3*, *arf6-2* and *arf8-3* mutants. Mutant alleles are homozygous unless specified as heterozygous, including arf6+/- and arf8+/-. **b-c**, Epistasis analysis of *arf6, nph4-1* and *arf8*. The *nph4-1* mutation did not display parthenocarpy after emasculation. In **b**, photo showing representative pistils of different genotypes 7 days after emasculation. Bar = 2 mm. In **c**, pistil lengths, n>15. In boxplots **a** and **c**, center lines and box edges are medians and the lower/upper quartiles, respectively. Whiskers extend to the lowest and highest data points within 1.5x interquartile range (IQR) below and above the lower and upper quartiles, respectively. Different letters above the boxes represent significant differences ($p < 0.05$) as determined by Tukey's HSD mean separation test. In **a-c**, two biological repeats showed similar results.

Supplementary Figure 3

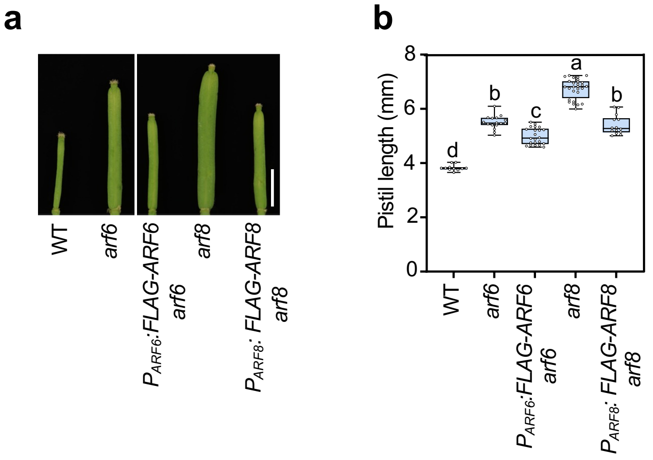


**Supplementary Figure 3. FLAG-ARF6/8 partially rescued the *arf6* and *arf8* parthenocarpic phenotype. a,** photo showing representative pistils of different lines 7 days after emasculation. Bar = 2 mm. **b**, pistil lengths. n>15. The center lines and box edges in the box plot are medians and the lower/upper quartiles, respectively. Whiskers extend to the lowest and highest data points within 1.5x IQR below and above the lower and upper quartiles, respectively. Different letters above the boxes represent significant differences ($p < 0.05$) as determined by Tukey's HSD mean separation test. Two biological repeats showed similar results.

Supplementary Figure 4

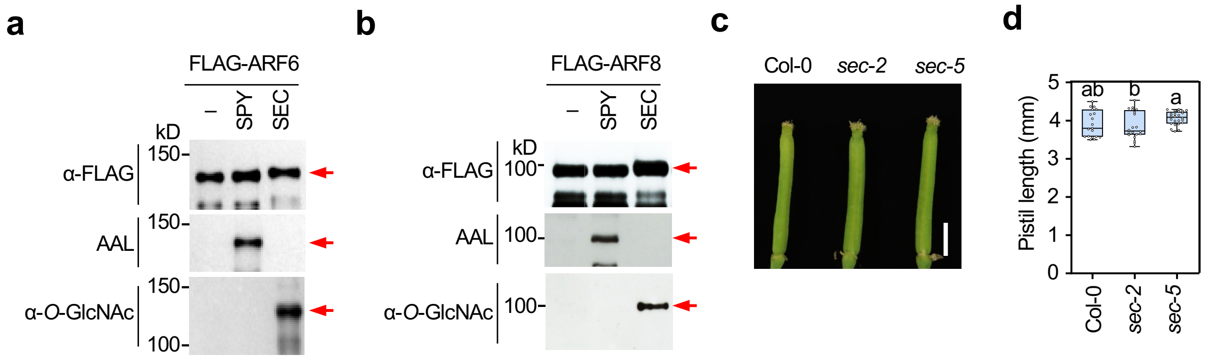


**Supplementary Figure 4. ARF6 and ARF8 are *O*-fucosylated and *O*-GlcNAcylated. a-b,** FLAG-ARF6 and -ARF8 were *O*-fucosylated by SPY and *O*-GlcNAcylated by SEC, respectively. FLAG-ARF6/8 proteins were expressed alone or co-expressed with SPY or SEC in *N. benthamiana*. Immunoblots containing affinity-purified FLAG-ARF6/8 were probed with anti-FLAG, AAL-biotin or anti-*O*-GlcNAc antibody as labeled. Arrow in the top panel indicates FLAG-ARF6 or -ARF8, arrow in the middle panel indicates *O*-fucosylated FLAG-ARF6 or -ARF8, and arrow in the bottom panel indicates *O*-GlcNAcylated FLAG-ARF6 or -ARF8. **c-d,** *sec* mutants did not alter growth of unpollinated pistils. **c,** photo showing representative pistils of different genotypes 7 days after emasculation. Bar = 1 mm. **d**, pistil lengths. n>15. The center lines and box edges in the box plot are medians and the lower/upper quartiles, respectively. Whiskers extend to the lowest and highest data points within 1.5x IQR below and above the lower and upper quartiles, respectively. Different letters above the boxes represent significant differences ($p < 0.05$) as determined by Tukey's HSD mean separation test. Two biological repeats showed similar results.

Supplementary Figure 5

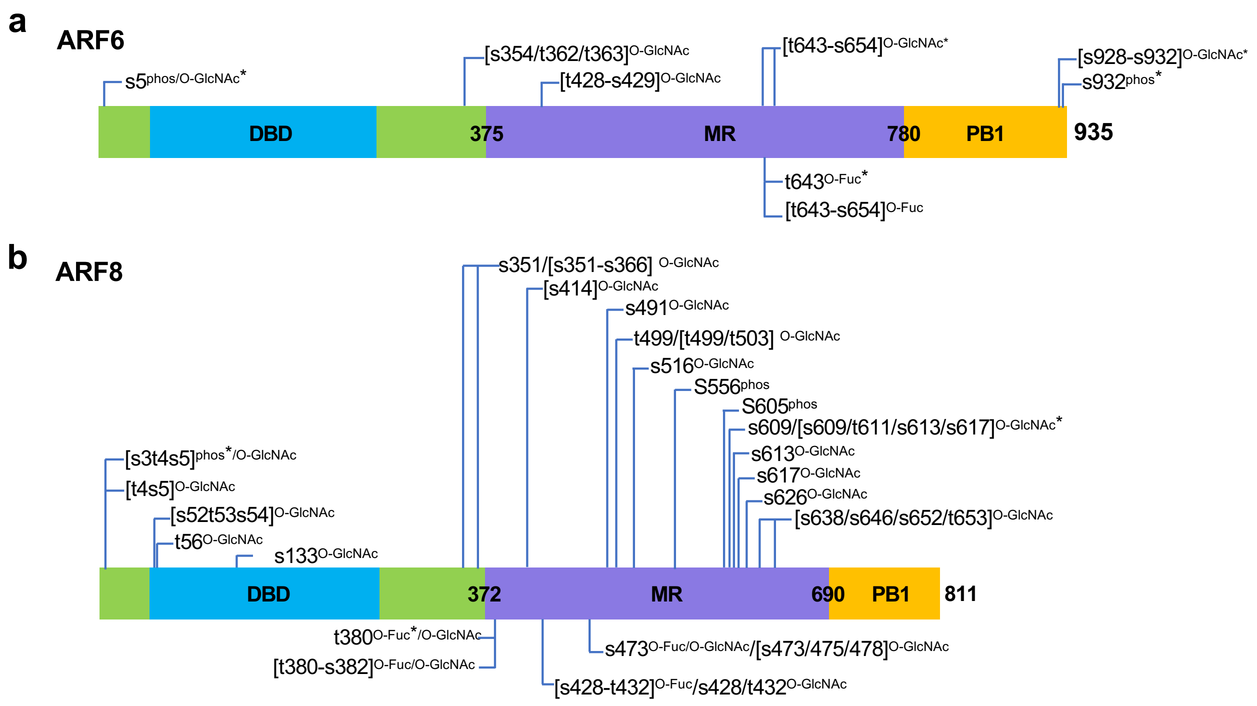


**Supplementary Figure 5. PTM sites in ARF6 and ARF8. a-b,** *O*-Fuc, *O*-GlcNAc and phosphorylation sites in ARF6 and ARF8 identified by MS analysis. The schematic shows the ARF6 (**a**) or ARF8 protein (**b**); The marked S/T residues are confirmed PTM sites. The sequence within brackets contains PTM(s) for which the specific residue(s) could not be determined. * indicates PTM that was reported previously[35,47].

Supplementary Figure 6

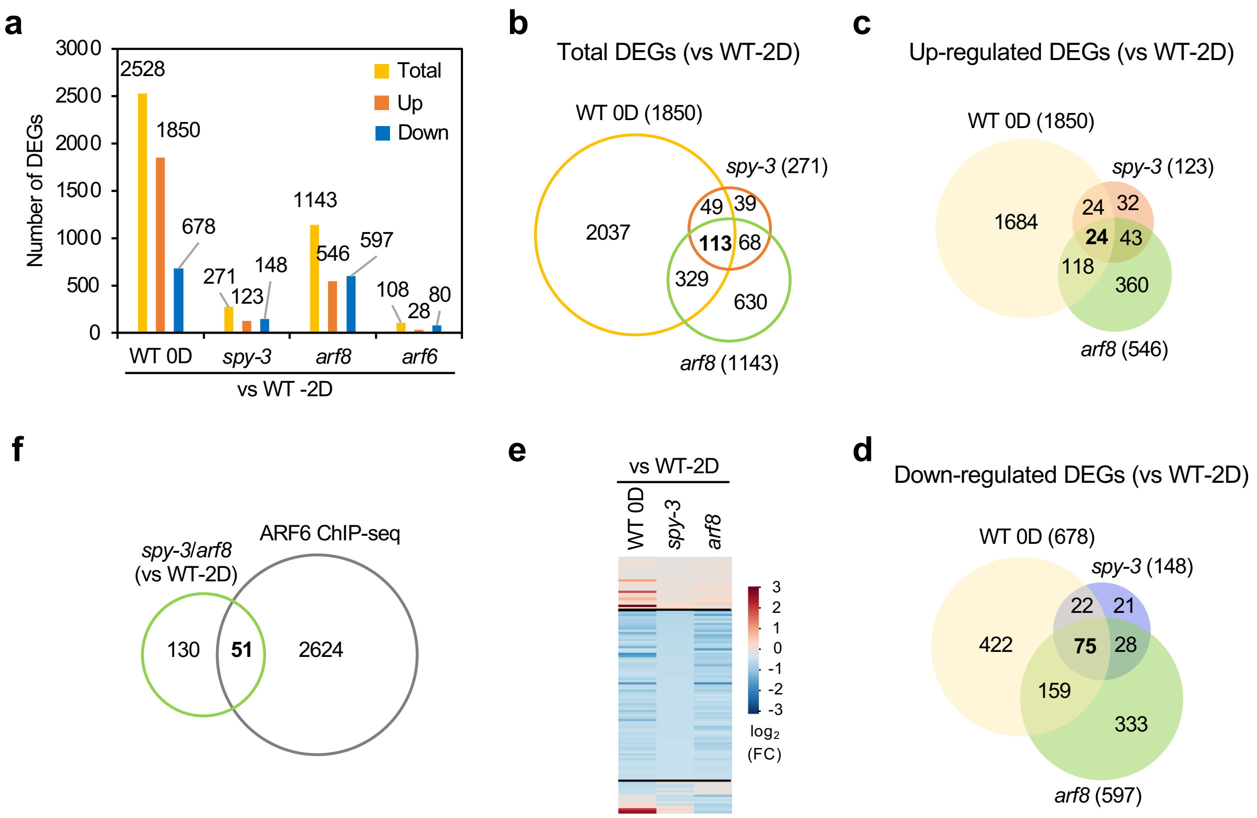


**Supplementary Figure 6. Identification of coregulated genes among fertilization-responsive vs ARF6-, ARF8- and SPY-responsive genes in pistils by RNA-seq analysis.** RNA-seq analysis was performed using −2 DAA pistils of *arf6, arf8, spy-3* and WT, and 0 DAA WT pistils. The differentially expressed gene (DEG) lists for ARF6-, ARF8- and SPY-responsive genes, and for fertilization-responsive genes (WT 0 DAA vs WT –2 DAA) are in **Supplementary Table 2**. **a,** Total, up- or down-regulated DEGs in WT 0 DAA or in each mutant (vs WT –2 DAA). **b-d.** Venn diagrams of coregulated DEGs among fertilization (WT 0 DAA vs WT –2 DAA), *arf6, arf8* and *spy-3*. **e**, Heat map of coregulated genes among fertilization-, ARF8- and SPY-responsive DEGs. **f**, Venn diagram of overlapping genes between total SPY/ARF8 coregulated genes (181 DEGs) and the ARF6 ChIP-seq gene list[51].

### Supplementary Figure 7

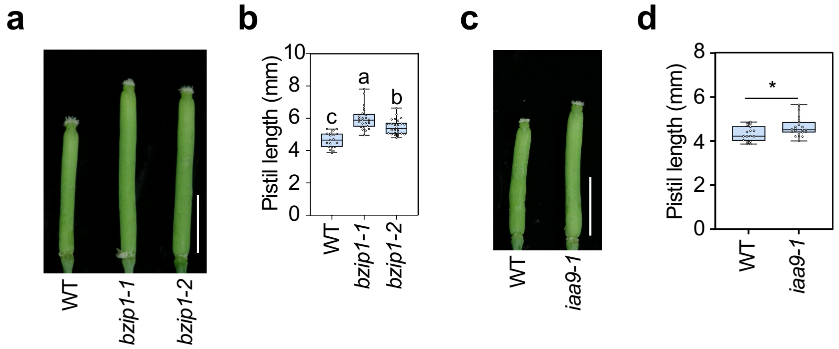


**Supplementary Figure 7. The *bzip1* and *iaa9* mutants displayed longer pistils after emasculation.** In **a** and **c**, photos showing representative pistils of different genotypes 7 days after emasculation. Bar = 2 mm. In **b** and **d**, pistil lengths were shown in boxplots. n>15. The center lines and box edges are medians and the lower/upper quartiles, respectively. Whiskers extend to the lowest and highest data points within 1.5x IQR below and above the lower and upper quartiles, respectively. Different letters (in **b**) or the asterisk (in **d**) above the boxes represent significant differences ($p < 0.05$) as determined by Tukey's HSD mean separation test. Two biological repeats showed similar results.

Supplementary Figure 8

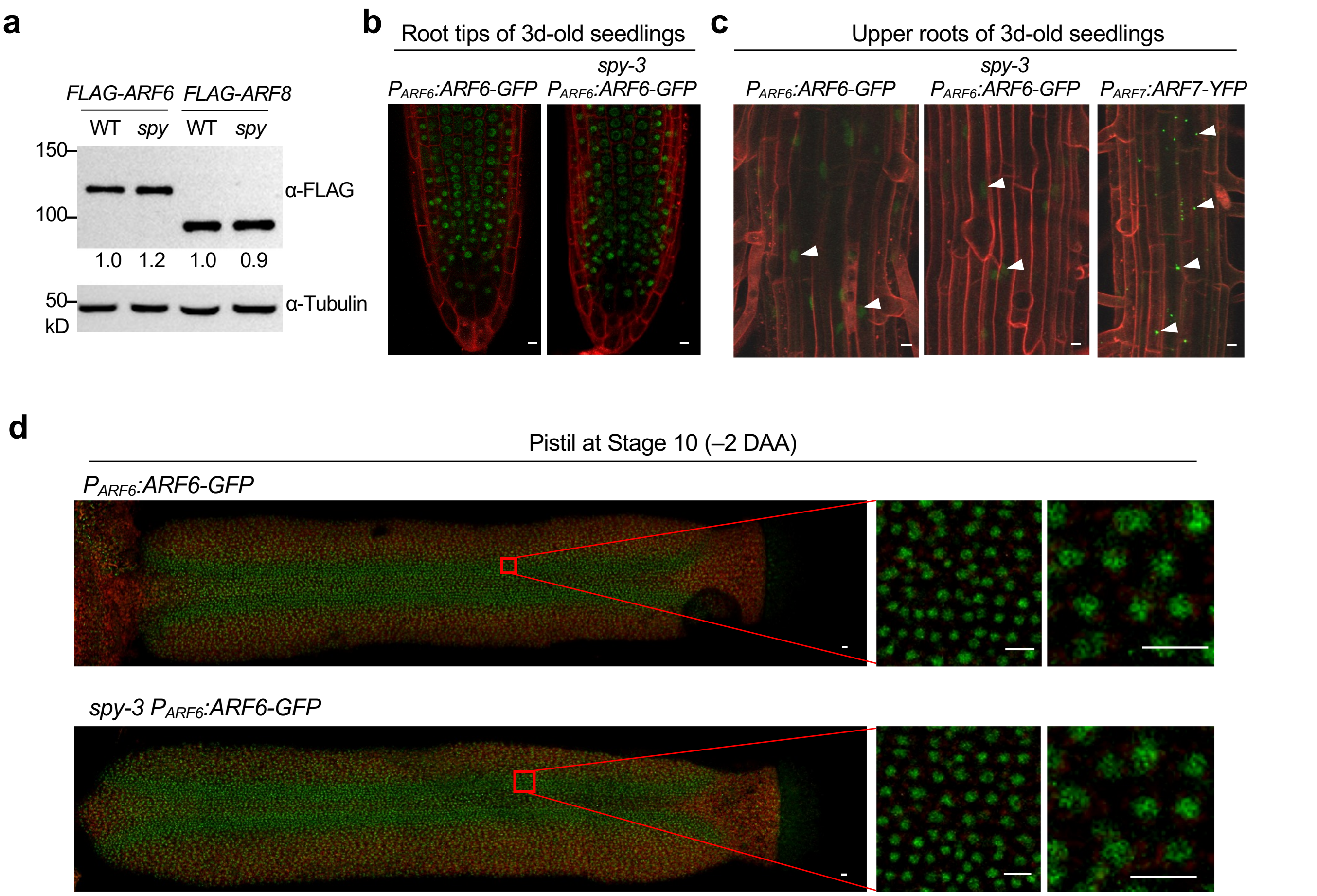


**Supplementary Figure 8. ARF6 and ARF8 protein accumulation or nuclear localization were not affected by *spy*. a**, $P_{UBQ10}$:*FLAG-ARF6* and $P_{UBQ10}$:*FLAG-ARF8* in WT vs *spy* background. Immunoblots containing total proteins extracted from seedlings were probed with anti-FLAG or anti-tubulin antibody. **b-d**, $P_{ARF6}$:*ARF6-GFP* in WT vs *spy* background. In **c**, $P_{ARF7}$:*ARF7-YFP* in WT background was included as a control. GFP/YFP signals detected by confocal microscopy showing root tips (**b**) or upper roots (**c**) of 3d-old seedlings or stage-10 pistils (**d**). In **c,** ARF6-GFP was only detected in the nuclei of root cells in the maturation zone, whereas ARF7-YFP localized in cytoplasmic condensates. In **b-c**, roots were stained with propidium iodide before imaging. In **b-d**, bar = 10 μm.

Supplementary Figure 9

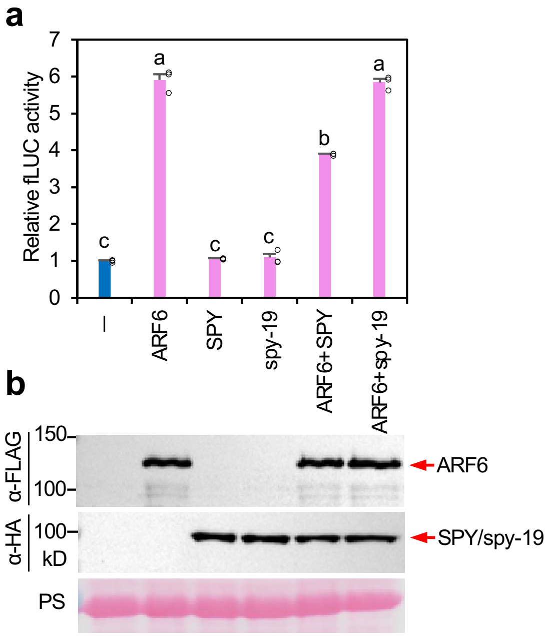


**Supplementary Figure 9. ARF6 transactivation activities were reduced by SPY, but not by spy-19. a-b,** Dual luciferase assay in the *N. benthamiana* transient expression system. *35S:Renilla LUC (rLUC)* was the internal control for transformation efficiency. The reporter construct contained *P3(2x):fLUC*. Effector constructs included *35S:FLAG-ARF6* and/or *35S:HA-SPY,* and/or *35S:HA-spy-19* as labeled. In **a**, relative fLUC activity was calculated by normalizing with rLUC activity in each sample. Means ± SE of 3 biological replicas are shown. Different letters above the bars represent significant differences ($p < 0.05$) as determined by Tukey's HSD mean separation test. In **b,** each effector protein was expressed at similar levels in different samples. Effector proteins in *N. benthamiana* extracts were detected by immunoblot using anti-FLAG, anti-Myc and anti-HA antibodies as labeled. Ponceau S (PS)-stained gel images showing similar sample loading. Two biological repeats showed similar results.

Supplementary Fig. 10

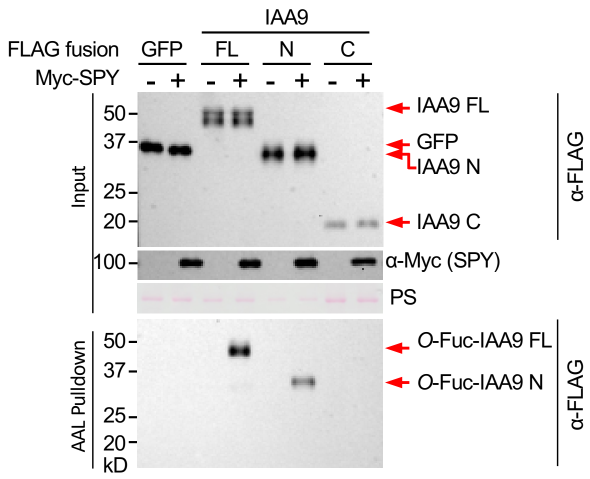


**Supplementary Figure 10. AAL pulldown assay showed that N-terminal domain of IAA9 was *O*-fucosylated by SPY.** FLAG-tagged full-length (FL) or truncated IAA9 proteins were expressed alone (–) or co-expressed (+) with Myc-SPY in *N. benthamiana.* FLAG-GFP*,* a negative control. *O*-fucosylated proteins were pull-downed by AAL-agarose. Immunoblot containing input (top panel) or AAL-agarose pull-down samples (bottom panel) was probed with anti-FLAG and anti-Myc antibodies as labeled. PS, Ponceau S-stained blot showing even loading. N, N-terminal domain of IAA9 (amino acid residues 1-208); C, C-terminal PB1 domain of IAA9 (amino acid residues 209-326). Two biological repeats showed similar results.

Supplementary Fig. 11

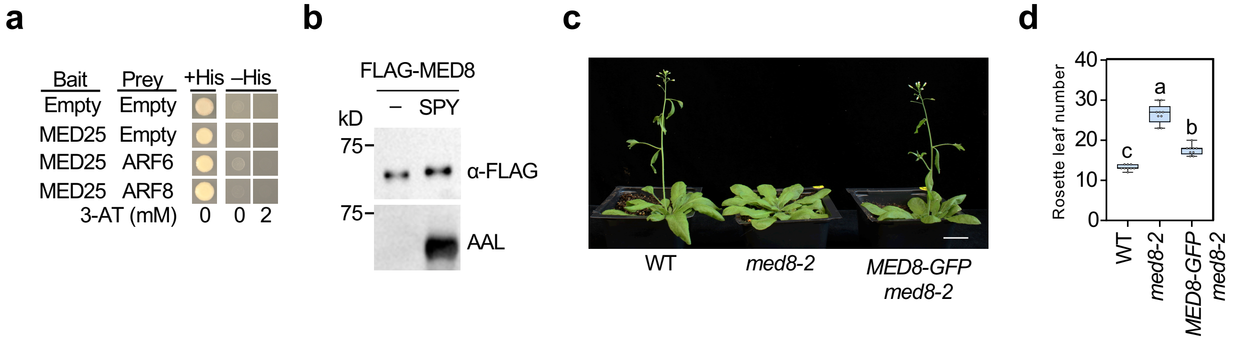


**Supplementary Figure 11. SPY *O*-fucosylates MED8. a,** ARF6/8 did not interact with MED25 in Y2H assay. **b,** MED8 was *O*-fucosylated by SPY. FLAG-MED8 was expressed alone (–) or co-expressed (+) with SPY in *N. benthamiana*. Immunoblots containing affinity-purified FLAG-MED8 proteins were probed with AAL-biotin or anti-FLAG antibody as labeled. **c-d,** The *35S:MED8-GFP* transgene rescued the late flowering phenotype of *med8-2*. In **c**, photo was taken at 30d-old, and bar = 2 cm. In **d**, n=10. The center lines and box edges in the box plot are medians and the lower/upper quartiles, respectively. Whiskers extend to the lowest and highest data points within 1.5x IQR below and above the lower and upper quartiles, respectively. Different letters above the boxes represent significant differences ($p < 0.05$) as determined by Tukey's HSD mean separation test. Two biological repeats showed similar results.
