## Supplementary Data Sets 1-4 for "The *O*-Fucosyltransferase SPINDLY Attenuates Auxin-Induced Fruit Growth by Inhibiting ARF6 and ARF8 binding to Coactivator Mediator Complex in *Arabidopsis*"

**Supplementary Data Set S1. ETD MS$^2$ spectrum of the tryptic ARF6 peptide T$^{O\text{-}Fuc}$NSAMTSSGWPSK from *N. Benthamiana.*** Precursor m/z = 750.3366. The *O*-Fuc-modification appears to be located on T1.

ETD Spectrum

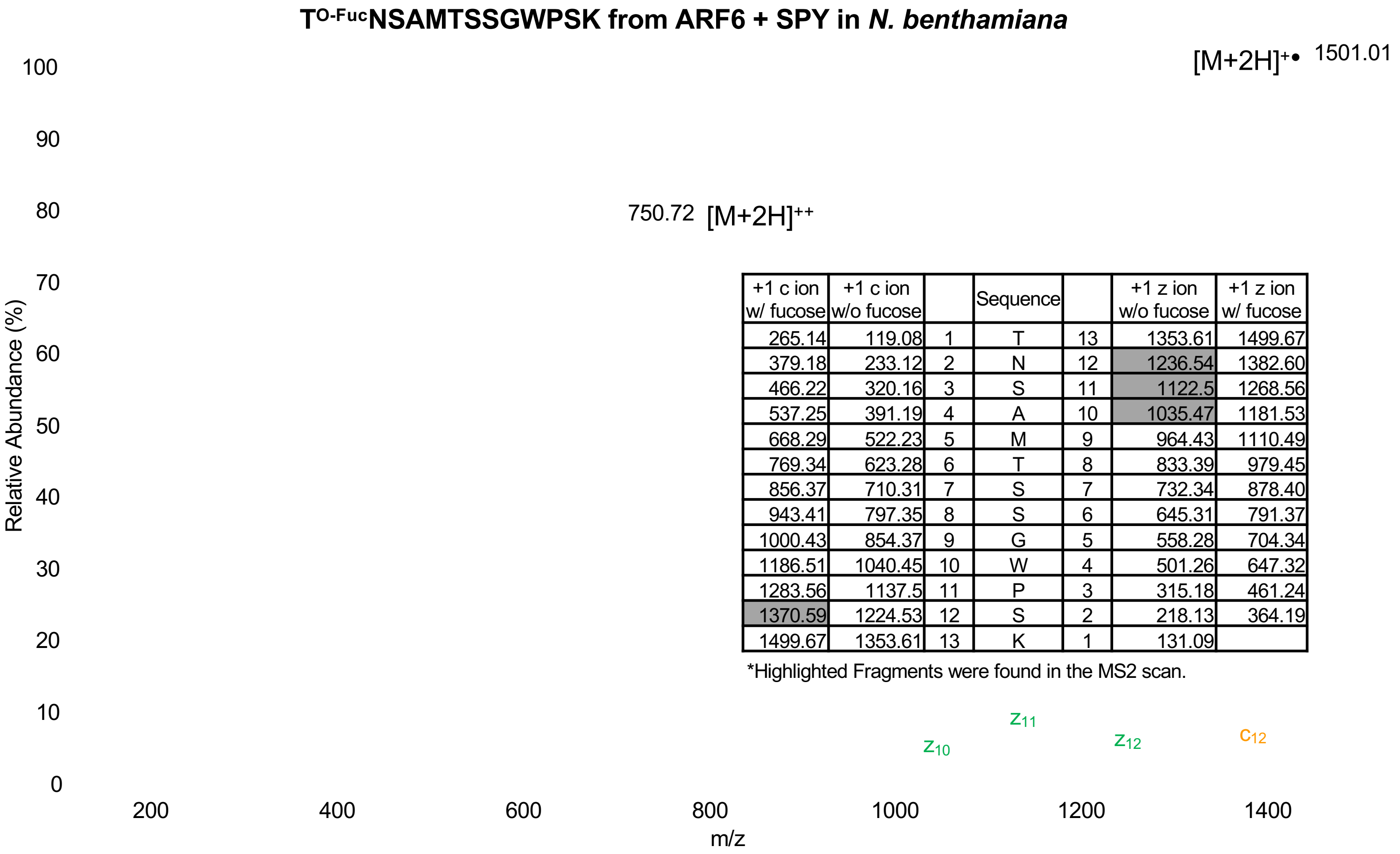


| +1 c ion w/ fucose | +1 c ion w/o fucose | | Sequence | | +1 z ion w/o fucose | +1 z ion w/ fucose |
|---|---|---|---|---|---|---|
| 265.14 | 119.08 | 1 | T | 13 | 1353.61 | 1499.67 |
| 379.18 | 233.12 | 2 | N | 12 | 1236.54 | 1382.60 |
| 466.22 | 320.16 | 3 | S | 11 | 1122.5 | 1268.56 |
| 537.25 | 391.19 | 4 | A | 10 | 1035.47 | 1181.53 |
| 668.29 | 522.23 | 5 | M | 9 | 964.43 | 1110.49 |
| 769.34 | 623.28 | 6 | T | 8 | 833.39 | 979.45 |
| 856.37 | 710.31 | 7 | S | 7 | 732.34 | 878.40 |
| 943.41 | 797.35 | 8 | S | 6 | 645.31 | 791.37 |
| 1000.43 | 854.37 | 9 | G | 5 | 558.28 | 704.34 |
| 1186.51 | 1040.45 | 10 | W | 4 | 501.26 | 647.32 |
| 1283.56 | 1137.5 | 11 | P | 3 | 315.18 | 461.24 |
| 1370.59 | 1224.53 | 12 | S | 2 | 218.13 | 364.19 |
| 1499.67 | 1353.61 | 13 | K | 1 | 131.09 | |

*Highlighted Fragments were found in the MS2 scan.

**Supplementary Data Set S2. ETD MS² spectrum of the tryptic ARF8 peptide RPWHAGT$^{O\text{-}Fuc}$SSLPDGR from *N. Benthamiana*.** Precursor m/z = 561.6119. The *O*-Fuc-modification appears to be located on T7.

ETD Spectrum

**RPWHAGT$^{O\text{-}Fuc}$SSLPDGR from ARF8 + SPY in *N. benthamiana***

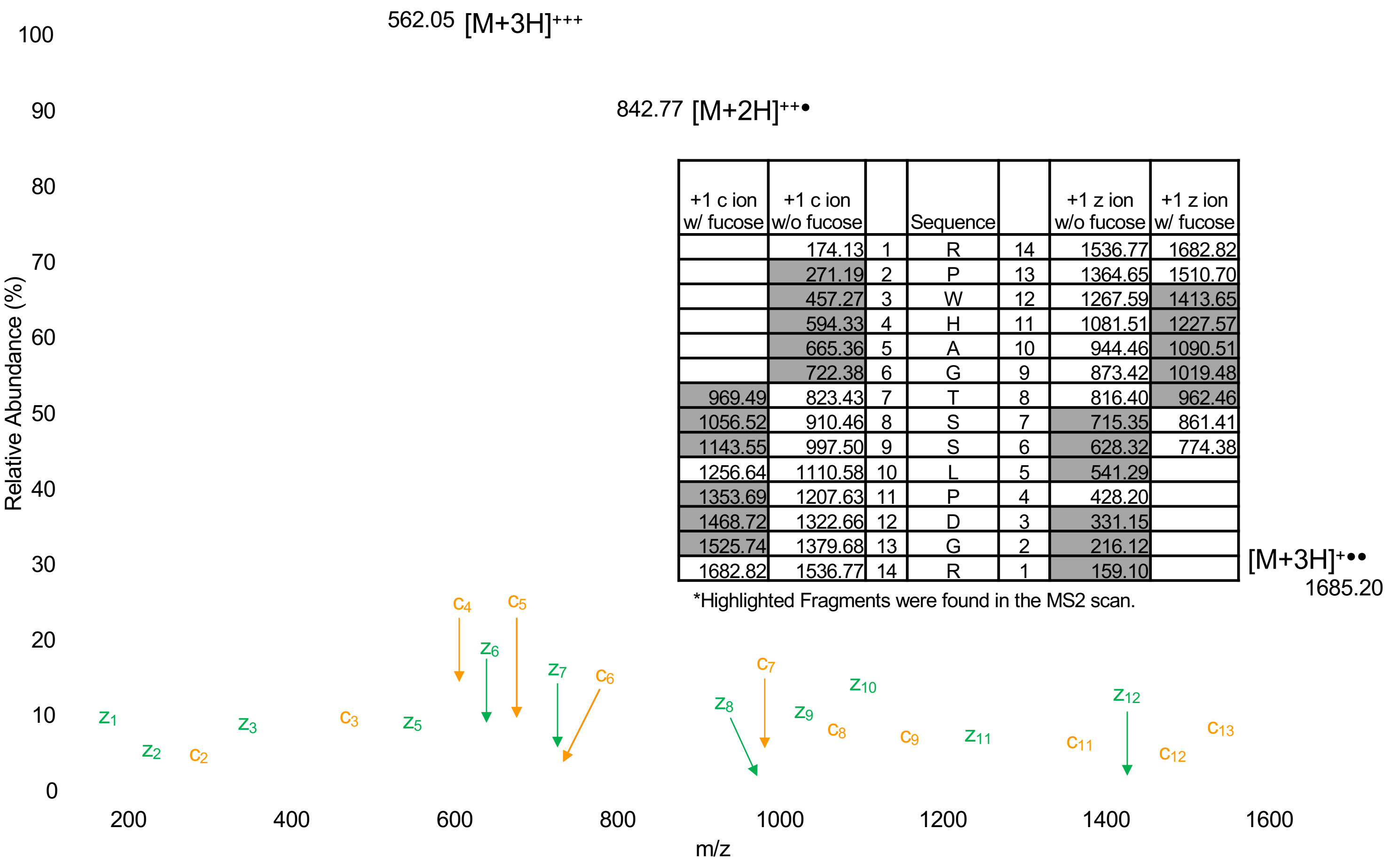


| +1 c ion w/ fucose | +1 c ion w/o fucose | | Sequence | | +1 z ion w/o fucose | +1 z ion w/ fucose |
|---|---|---|---|---|---|---|
| | 174.13 | 1 | R | 14 | 1536.77 | 1682.82 |
| | 271.19 | 2 | P | 13 | 1364.65 | 1510.70 |
| | 457.27 | 3 | W | 12 | 1267.59 | 1413.65 |
| | 594.33 | 4 | H | 11 | 1081.51 | 1227.57 |
| | 665.36 | 5 | A | 10 | 944.46 | 1090.51 |
| | 722.38 | 6 | G | 9 | 873.42 | 1019.48 |
| 969.49 | 823.43 | 7 | T | 8 | 816.40 | 962.46 |
| 1056.52 | 910.46 | 8 | S | 7 | 715.35 | 861.41 |
| 1143.55 | 997.50 | 9 | S | 6 | 628.32 | 774.38 |
| 1256.64 | 1110.58 | 10 | L | 5 | 541.29 | |
| 1353.69 | 1207.63 | 11 | P | 4 | 428.20 | |
| 1468.72 | 1322.66 | 12 | D | 3 | 331.15 | |
| 1525.74 | 1379.68 | 13 | G | 2 | 216.12 | |
| 1682.82 | 1536.77 | 14 | R | 1 | 159.10 | |

*Highlighted Fragments were found in the MS2 scan.

**Supplementary Data Set S3. HCD MS² spectrum of the tryptic ARF6 peptide LDL[SQMGT]$^{O\text{-}Fuc}$DNNQQ YQAMLAAGLQNIGGGDPLR from *N. Benthamiana*.** Precursor m/z =1179.5625. The HCD spectrum confirms the sequence of this peptide, but does not provide *O*-Fuc site information.

HCD Spectrum

**LDL[SQMGT]$^{O\text{-}Fuc}$DNNQQYQAMLAAGLQNIGGGDPLR from ARF8 + SPY in *N. benthamiana***

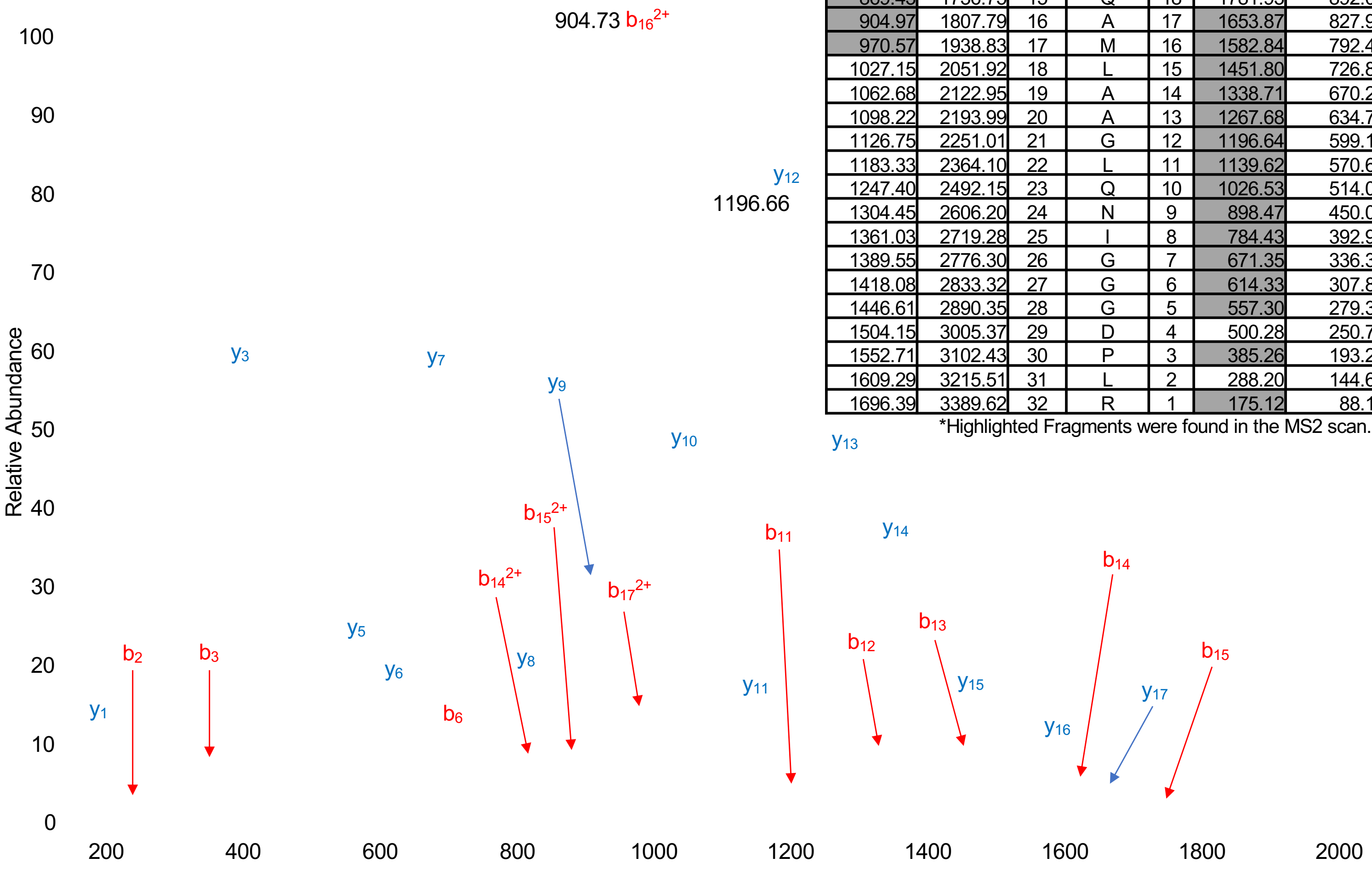


| +2 b ion w/o fucose | +1 b ion w/o fucose | | Sequence | | +1 y ion w/o fucose | +2 y ion w/o fucose |
|---|---|---|---|---|---|---|
| 57.59 | 114.09 | 1 | L | 32 | 3389.62 | 1696.39 |
| 115.13 | 229.12 | 2 | D | 31 | 3276.54 | 1639.81 |
| 171.71 | 342.20 | 3 | L | 30 | 3161.51 | 1582.27 |
| 215.25 | 429.23 | 4 | S | 29 | 3048.43 | 1525.69 |
| 279.32 | 557.29 | 5 | Q | 28 | 2961.39 | 1482.15 |
| 344.92 | 688.33 | 6 | M | 27 | 2833.34 | 1418.08 |
| 373.44 | 745.35 | 7 | G | 26 | 2702.30 | 1352.48 |
| 423.99 | 846.40 | 8 | T | 25 | 2645.27 | 1323.96 |
| 481.54 | 961.43 | 9 | D | 24 | 2544.23 | 1273.40 |
| 538.59 | 1075.47 | 10 | N | 23 | 2429.20 | 1215.86 |
| 595.64 | 1189.52 | 11 | N | 22 | 2315.16 | 1158.81 |
| 659.71 | 1317.57 | 12 | Q | 21 | 2201.11 | 1101.76 |
| 723.77 | 1445.63 | 13 | Q | 20 | 2073.05 | 1037.69 |
| 805.36 | 1608.70 | 14 | Y | 19 | 1945.00 | 973.62 |
| 869.43 | 1736.75 | 15 | Q | 18 | 1781.93 | 892.04 |
| 904.97 | 1807.79 | 16 | A | 17 | 1653.87 | 827.97 |
| 970.57 | 1938.83 | 17 | M | 16 | 1582.84 | 792.43 |
| 1027.15 | 2051.92 | 18 | L | 15 | 1451.80 | 726.83 |
| 1062.68 | 2122.95 | 19 | A | 14 | 1338.71 | 670.25 |
| 1098.22 | 2193.99 | 20 | A | 13 | 1267.68 | 634.71 |
| 1126.75 | 2251.01 | 21 | G | 12 | 1196.64 | 599.17 |
| 1183.33 | 2364.10 | 22 | L | 11 | 1139.62 | 570.65 |
| 1247.40 | 2492.15 | 23 | Q | 10 | 1026.53 | 514.07 |
| 1304.45 | 2606.20 | 24 | N | 9 | 898.47 | 450.00 |
| 1361.03 | 2719.28 | 25 | I | 8 | 784.43 | 392.95 |
| 1389.55 | 2776.30 | 26 | G | 7 | 671.35 | 336.37 |
| 1418.08 | 2833.32 | 27 | G | 6 | 614.33 | 307.84 |
| 1446.61 | 2890.35 | 28 | G | 5 | 557.30 | 279.32 |
| 1504.15 | 3005.37 | 29 | D | 4 | 500.28 | 250.79 |
| 1552.71 | 3102.43 | 30 | P | 3 | 385.26 | 193.25 |
| 1609.29 | 3215.51 | 31 | L | 2 | 288.20 | 144.69 |
| 1696.39 | 3389.62 | 32 | R | 1 | 175.12 | 88.11 |

*Highlighted Fragments were found in the MS2 scan.

**Supplementary Data Set S4. ETD MS$^2$ spectrum of the tryptic ARF8 peptide RPWHAGT$^{O\text{-}Fuc}$SSLPDGR from *N. benthamiana*.** Precursor m/z = 893.8301. The *O*-Fuc-modification appears to be located on S17.

ETD Spectrum **QQFVQLQEPHHQYLQQS$^{O\text{-}Fuc}$ASHNSDLMLQQQQQQASR from ARF8 + SPY in *N. benthamiana***

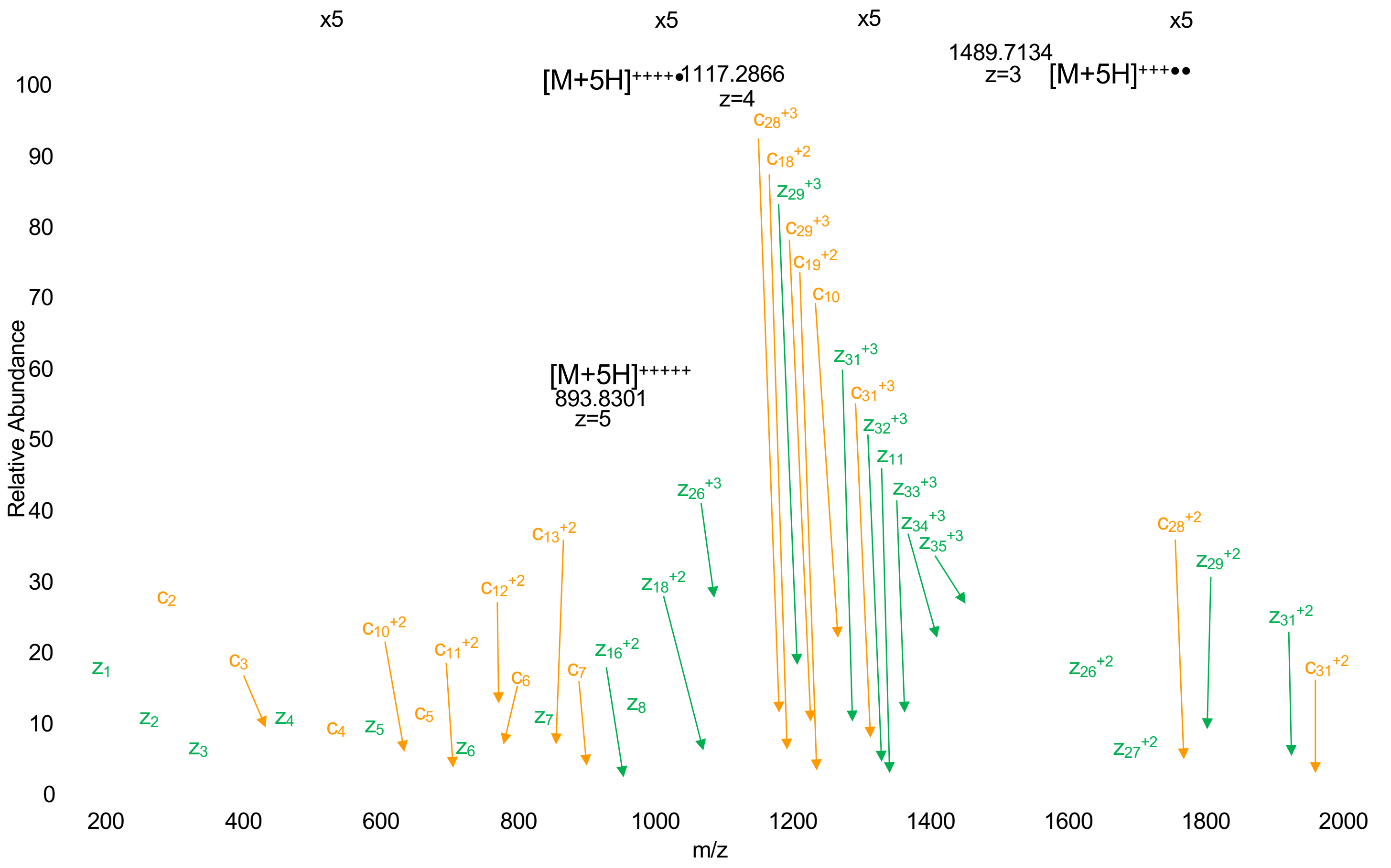


| +3 c ion w/ fucose | +3 c ion w/o fucose | +2 c ion w/ fucose | +2 c ion w/o fucose | +1 c ion w/ fucose | +1 c ion w/o fucose | | Sequence | | +1 z ion w/o fucose | +1 z ion w/ fucose | +2 z ion w/o fucose | +2 z ion w/ fucose | +3 z ion w/o fucose | +3 z ion w/ fucose |
|---|---|---|---|---|---|---|---|---|---|---|---|---|---|---|
| | 49.37 | | 73.55 | | 146.09 | 1 | Q | 36 | 4317.06 | 4463.12 | 2159.03 | 2232.06 | 1439.69 | 1488.38 |
| | 92.06 | | 137.58 | | 274.15 | 2 | Q | 35 | 4172.98 | 4319.04 | 2087.00 | 2160.02 | 1391.67 | 1440.35 |
| | 141.08 | | 211.11 | | 421.22 | 3 | F | 34 | 4044.92 | 4190.98 | 2022.97 | 2095.99 | 1348.98 | 1397.67 |
| | 174.10 | | 260.65 | | 520.29 | 4 | V | 33 | 3897.86 | 4043.91 | 1949.43 | 2022.46 | 1299.96 | 1348.64 |
| | 216.79 | | 324.68 | | 648.35 | 5 | Q | 32 | 3798.79 | 3944.85 | 1899.90 | 1972.93 | 1266.93 | 1315.62 |
| | 254.48 | | 381.22 | | 761.43 | 6 | L | 31 | 3670.73 | 3816.79 | 1835.87 | 1908.90 | 1224.25 | 1272.93 |
| | 297.17 | | 445.25 | | 889.49 | 7 | Q | 30 | 3557.64 | 3703.70 | 1779.33 | 1852.36 | 1186.55 | 1235.24 |
| | 340.18 | | 509.77 | | 1018.53 | 8 | E | 29 | 3429.59 | 3575.64 | 1715.30 | 1788.33 | 1143.87 | 1192.55 |
| | 372.53 | | 558.30 | | 1115.58 | 9 | P | 28 | 3300.54 | 3446.60 | 1650.78 | 1723.80 | 1100.85 | 1149.54 |
| | 418.22 | | 626.83 | | 1252.64 | 10 | H | 27 | 3203.49 | 3349.55 | 1602.25 | 1675.28 | 1068.50 | 1117.19 |
| | 463.91 | | 695.35 | | 1389.70 | 11 | H | 26 | 3066.43 | 3212.49 | 1533.72 | 1606.75 | 1022.82 | 1071.50 |
| | 506.59 | | 759.38 | | 1517.76 | 12 | Q | 25 | 2929.37 | 3075.43 | 1465.19 | 1538.22 | 977.13 | 1025.82 |
| | 560.95 | | 840.92 | | 1680.82 | 13 | Y | 24 | 2801.31 | 2947.37 | 1401.16 | 1474.19 | 934.44 | 983.13 |
| | 598.64 | | 897.46 | | 1793.91 | 14 | L | 23 | 2638.25 | 2784.31 | 1319.63 | 1392.66 | 880.09 | 928.77 |
| | 641.33 | | 961.49 | | 1921.97 | 15 | Q | 22 | 2525.17 | 2671.23 | 1263.09 | 1336.12 | 842.39 | 891.08 |
| | 684.01 | | 1025.52 | | 2050.03 | 16 | Q | 21 | 2397.11 | 2543.17 | 1199.06 | 1272.09 | 799.71 | 848.39 |
| 761.71 | 713.02 | 1142.06 | 1069.03 | 2283.12 | 2137.06 | 17 | S | 20 | 2269.05 | 2415.11 | 1135.03 | 1208.06 | 757.02 | 805.71 |
| 785.39 | 736.70 | 1177.58 | 1104.55 | 2354.15 | 2208.09 | 18 | A | 19 | 2182.02 | 2328.08 | 1091.51 | 1164.54 | 728.01 | 776.70 |
| 814.40 | 765.71 | 1221.10 | 1148.07 | 2441.18 | 2295.13 | 19 | S | 18 | 2110.98 | 2257.04 | 1055.99 | 1129.02 | 704.33 | 753.02 |
| 860.09 | 811.40 | 1289.63 | 1216.60 | 2578.24 | 2432.19 | 20 | H | 17 | 2023.95 | 2170.01 | 1012.48 | 1085.51 | 675.32 | 724.01 |
| 898.10 | 849.41 | 1346.65 | 1273.62 | 2692.29 | 2546.23 | 21 | N | 16 | 1886.89 | 2032.95 | 943.95 | 1016.98 | 629.63 | 678.32 |
| 927.11 | 878.43 | 1390.16 | 1317.13 | 2779.32 | 2633.26 | 22 | S | 15 | 1772.85 | 1918.90 | 886.93 | 959.96 | 591.62 | 640.31 |
| 965.45 | 916.77 | 1447.68 | 1374.65 | 2894.35 | 2748.29 | 23 | D | 14 | 1685.82 | 1831.87 | 843.41 | 916.44 | 562.61 | 611.30 |
| 1003.15 | 954.46 | 1504.22 | 1431.19 | 3007.43 | 2861.37 | 24 | L | 13 | 1570.79 | 1716.85 | 785.90 | 858.93 | 524.27 | 572.95 |
| 1046.83 | 998.14 | 1569.74 | 1496.71 | 3138.47 | 2992.41 | 25 | M | 12 | 1457.70 | 1603.76 | 729.36 | 802.38 | 486.57 | 535.26 |
| 1084.52 | 1035.84 | 1626.28 | 1553.25 | 3251.55 | 3105.50 | 26 | L | 11 | 1326.66 | 1472.72 | 663.84 | 736.86 | 442.89 | 491.58 |
| 1127.21 | 1078.52 | 1690.31 | 1617.28 | 3379.61 | 3233.55 | 27 | Q | 10 | 1213.58 | 1359.64 | 607.29 | 680.32 | 405.20 | 453.88 |
| 1169.90 | 1121.21 | 1754.34 | 1681.31 | 3507.67 | 3361.61 | 28 | Q | 9 | 1085.52 | 1231.58 | 543.26 | 616.29 | 362.51 | 411.20 |
| 1212.58 | 1163.90 | 1818.37 | 1745.34 | 3635.73 | 3489.67 | 29 | Q | 8 | 957.46 | 1103.52 | 479.23 | 552.26 | 319.83 | 368.51 |
| 1255.27 | 1206.58 | 1882.40 | 1809.37 | 3763.79 | 3617.73 | 30 | Q | 7 | 829.40 | 975.46 | 415.21 | 488.23 | 277.14 | 325.83 |
| 1297.95 | 1249.27 | 1946.43 | 1873.40 | 3891.85 | 3745.79 | 31 | Q | 6 | 701.35 | 847.40 | 351.18 | 424.21 | 234.45 | 283.14 |
| 1340.64 | 1291.95 | 2010.46 | 1937.43 | 4019.91 | 3873.85 | 32 | Q | 5 | 573.29 | 719.34 | 287.15 | 360.18 | 191.77 | 240.45 |
| 1383.33 | 1334.64 | 2074.49 | 2001.46 | 4147.96 | 4001.91 | 33 | Q | 4 | 445.23 | 591.29 | 223.12 | 296.15 | 149.08 | 197.77 |
| 1407.01 | 1358.32 | 2110.00 | 2036.98 | 4219.00 | 4072.94 | 34 | A | 3 | 317.17 | 463.23 | 159.09 | 232.12 | 106.39 | 155.08 |
| 1436.02 | 1387.33 | 2153.52 | 2080.49 | 4306.03 | 4159.98 | 35 | S | 2 | 246.13 | 392.19 | 123.57 | 196.60 | 82.72 | 131.40 |
| 1488.38 | 1439.69 | 2232.06 | 2159.03 | 4463.12 | 4317.06 | 36 | R | 1 | 159.10 | | 80.05 | | 53.70 | |

*Highlighted Fragments were found in the MS2 scan.
